# Cell-free assays reveal that the HIV-1 capsid protects reverse transcripts from cGAS

**DOI:** 10.1101/2024.04.22.590513

**Authors:** Tiana M. Scott, Lydia M. Arnold, Jordan A. Powers, Delaney A. McCann, Ana B. Rowe, Devin E. Christensen, Miguel J. Pereira, Wen Zhou, Rachel M. Torrez, Janet H. Iwasa, Philip J. Kranzusch, Wesley I. Sundquist, Jarrod S. Johnson

**Author notes:** Contributed equally (Tiana M. Scott, Lydia M. Arnold).

## Abstract

Retroviruses can be detected by the innate immune sensor cyclic GMP-AMP synthase (cGAS), which recognizes reverse-transcribed DNA and activates an antiviral response.

However, the extent to which HIV-1 shields its genome from cGAS recognition remains unclear. To study this process in mechanistic detail, we reconstituted reverse transcription, genome release, and innate immune sensing of HIV-1 in a cell-free system. We found that wild-type HIV-1 capsids protect viral genomes from cGAS even after completing reverse transcription. Viral DNA could be “deprotected” by thermal stress, capsid mutations, or reduced concentrations of inositol hexakisphosphate (IP6) that destabilize the capsid. Strikingly, the capsid inhibitor lenacapavir also disrupted viral cores and dramatically potentiated cGAS activity, both in vitro and in cellular infections. Our results provide biochemical evidence that the HIV-1 capsid lattice conceals the genome from cGAS and that chemical or physical disruption of the viral core can expose HIV-1 DNA and activate innate immune signaling.

## Introduction

HIV-1 encapsidates the viral RNA genome and enzymes necessary for replication within a conical capsid comprising ∼1200 copies of the CA protein. Following viral fusion with a target cell membrane, the capsid and its contents (collectively known as the “core”) are released into the cytosol. This initiates reverse transcription, which converts two copies of the single-stranded RNA genome into one copy of double-stranded DNA (dsDNA). Reverse transcription proceeds inside an intact, or largely intact, capsid as the viral core is trafficked from the cell surface to an integration site in the nucleus^1–10^. During a successful infection, reverse transcription likely completes in the nucleus^4,11–14^, with capsid uncoating and genome release occurring only moments before viral integration into host chromatin^4,5^. Thus, the HIV-1 capsid is thought to shield the viral genome from cytosolic innate immune sensors during entry^15–17^. Yet, under some conditions, retroviral capsids fail to protect reverse transcribed viral DNA completely, permitting its detection in dendritic cells and macrophages and enabling the downstream induction of interferon (IFN) and inflammatory cytokines^16–21^. Mechanistic studies of HIV-1 sensing have remained challenging, however, because detection of reverse transcripts is a rare event that happens deep within these specialized cells.

Cyclic GMP-AMP synthase (cGAS) is the primary innate immune sensor that detects retroviral DNA^17,18^. Upon binding dsDNA in a sequence-independent manner, cGAS synthesizes the dinucleotide second messenger 2′3′-cyclic GMP-AMP (cGAMP), which binds and activates the adapter protein Stimulator of IFN genes (STING). STING then promotes downstream antiviral activities, including activation of the transcription factors IRF3 and NF-κB, which translocate to the nucleus and drive expression of type I and type III IFN^22,23^. IFN is rapidly secreted and initiates autocrine or paracrine signaling through IFN receptors. This upregulates hundreds of IFN-stimulated genes (ISGs), many of which can potently restrict HIV-1 infection^24–27^. Capsid structural integrity apparently influences the degree to which cGAS recognizes viral DNA, as specific CA mutations and CA-binding proteins are linked with increased or decreased IFN responses during infection^15–17,21,28–33^. However, the parameters that regulate exposure of viral DNA and its detection by cGAS remain unclear.

Here, we have established a cell-free system for studying the essential steps in cGAS-mediated sensing of HIV-1. We find that wild-type (wt) HIV-1 capsids remain stable and protect their genomes from cGAS detection, even several days after reverse transcription. We also identified conditions and drug treatments that “deprotected” viral genomes and increased cGAS activity in vitro and in cells. Thus, the HIV-1 capsid is a molecular cloak that shields the viral genome from innate immune sensing.

## Results

### Reconstitution of HIV-1 innate immune sensing in vitro

The discovery that HIV-1 viral capsids are stabilized by IP6 has facilitated biochemical studies of endogenous reverse transcription and host-virus interactions^3,34–36^. We sought to extend these methods to study the physical and biochemical principles that control cGAS-mediated detection of reverse transcripts. To perform efficient endogenous reverse transcription (ERT), we permeabilized the viral membrane using the pore-forming peptide melittin, stabilized viral capsids in buffers that contained IP6 and ribonucleotides at the physiological levels observed in macrophages, and provided dNTP substrates for reverse transcription (Fig. 1A).

**Figure 1.**
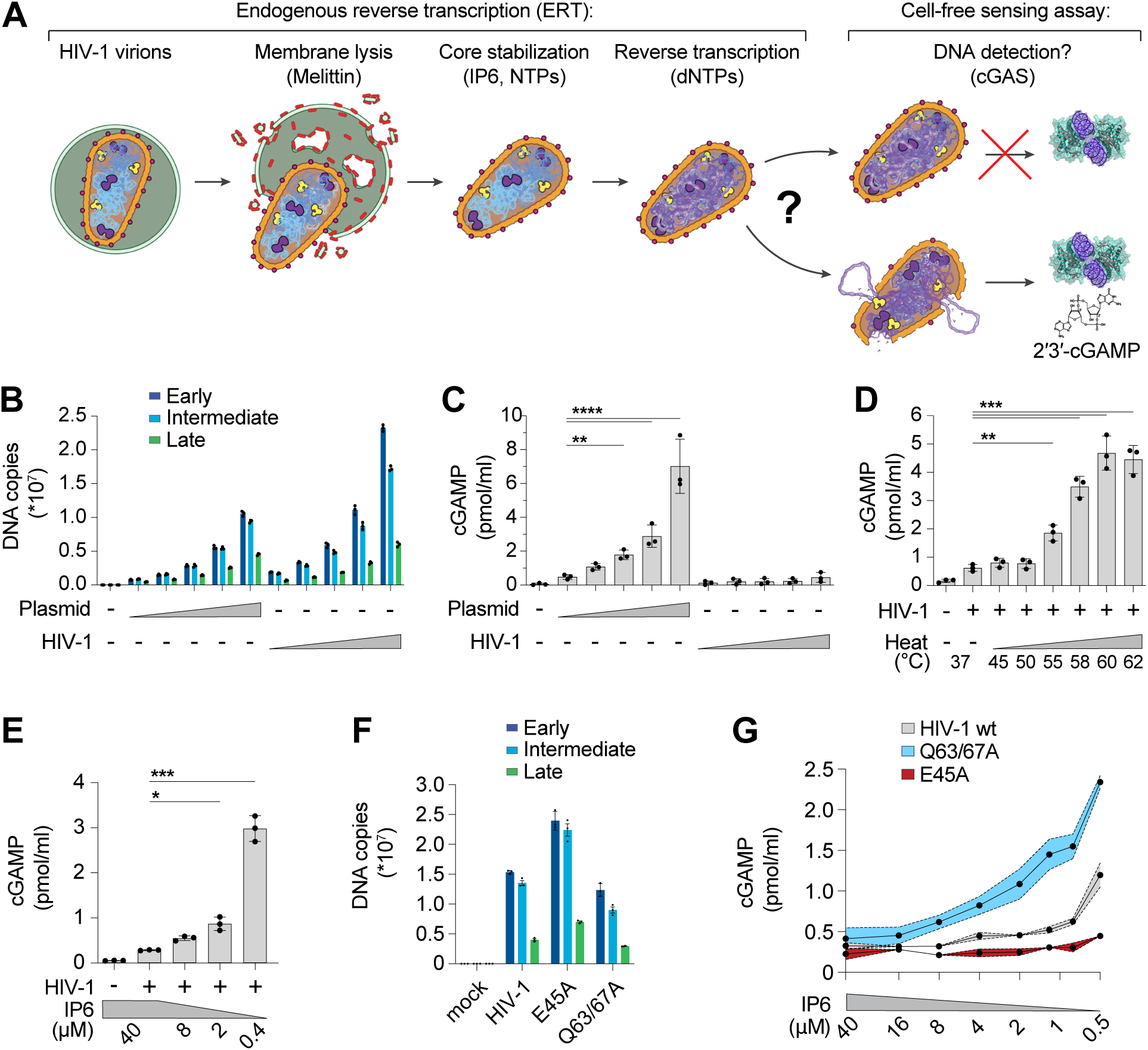
The HIV-1 capsid protects RT products from cGAS in vitro. **(A)** Steps in the endogenous reverse transcription (ERT) and cell-free sensing assays. Capsids that remain intact after ERT may protect viral DNA from detection and capsids that lose integrity may permit cGAS activation and cGAMP production. **(B)** qPCR measurements of plasmid DNA control containing the viral genome sequence compared to HIV-1 ERT reactions in 2-fold dilution series, incubated for 16 h under identical “standard” conditions at 37°C (including melittin, IP6, NTPs, dNTPs, MgCl_2_, NaCl, and BSA in a Tris buffer to approximate cytoplasmic conditions as stated in the Methods). Primer sets detect sequences for early (minus strand strong stop), intermediate (first strand transfer), and late RT (second strand transfer), which are present in both the plasmid and viral genome. **(C)** Cell-free sensing assay for samples shown in (B). After ERT, samples were incubated with recombinant cGAS for 8 h at 37°C. cGAMP levels were determined by ELISA (this applies to all cell-free sensing assays). **(D)** Cell-free sensing assay of heat treated ERT samples. Samples from standard ERT reactions (16 h at 37°C) were held at the indicated temperatures (20 s) and then subject to cell-free sensing assays performed as described in the Methods section. **(E)** Samples from standard ERT reactions (16 h at 37°C) performed with cellular levels of IP6 (40 µM) were diluted at equivalent amounts into cell-free sensing assay reactions that contained a range of IP6 (40 µM to 0.4 µM final concentration). **(F)** qPCR measurements of DNA copy numbers from ERT reactions with wt HIV-1 virions compared to virions with stabilizing (E45A) or destabilizing (Q63/67A) mutations in CA. **(G)** Cell-free sensing assay of samples from (F) after normalizing for DNA input and diluting into cGAS reactions at the indicated IP6 concentrations. Statistics were calculated using a one-way ANOVA with Tukey’s multiple comparisons test: p<0.05: *, p<0.01: **, p<0.001: ***, p<0.0001: ****. Graphs depict mean ± SD from three samples from a representative experiment, selected from three independent experiments.

Samples were then incubated with recombinant human cGAS to determine how efficiently the ERT products activated cGAS to produce cGAMP. cGAS activation levels were compared to those produced by a range of naked plasmid DNA concentrations. Relative DNA levels were measured by quantitative PCR (qPCR) detection of plasmid and full-length HIV-1_SG3_ genomes with primer sets that detected early, intermediate, and late reverse transcripts, and the equivalent DNA segments in the plasmid control (Fig. 1B). ERT was highly efficient, as previously reported^3^, yielding roughly 0.5 copies of late reverse transcripts per input viral core. Levels of early and intermediate products are higher because there are two early and intermediate primer binding sites on each copied genome, and because RT can initiate on both packaged RNA strands (before ultimately resolving to a single copy of dsDNA). As expected, plasmid DNA strongly activated cGAS in a concentration-dependent fashion. However, the HIV-1 ERT samples did not activate cGAS, suggesting that the viral DNA remained protected inside the capsid (Fig. 1C).

Performing ERT and cell-free sensing assays in two stages allowed us to uncouple parameters that influence ERT from parameters that impact cGAS activity (Fig. 1A). Viral cores that successfully completed RT were destabilized by limited heat treatment, which exposed the viral genome to cGAS (Fig. 1D). Under these conditions, the viral DNA was sensed efficiently. In contrast, heat treatment of plasmid control samples did not alter cGAS activity (S1 Fig. A-C).

Linear and circular forms of control DNA samples were detected at similar levels (S1 Fig. D-F), which agrees with previous findings using recombinant cGAS assays^37^, and indicates that linear viral DNA and control circular plasmid DNA should be sensed equivalently if the viral DNA were fully exposed. Together, these data reinforce the idea that viral DNA is protected by the capsid.

HIV capsid stability can also be modulated by IP6^34,35^ or by mutations in CA^3,38–42^. We therefore tested the effects of both variables on cGAS sensing of ERT products. Reducing IP6 from the cellular 40 µM concentration^43^ to 0.4 µM following reverse transcription increased cGAS detection of viral DNA by ∼10-fold (Fig. 1E). cGAS sensing was also altered by CA mutations that either stabilize (E45A) or destabilize (Q63/67A) capsids^3,38^. Specifically, we observed that the destabilizing Q63/67A mutant enhanced cGAS activation (normalized for cDNA levels), particularly as IP6 levels were reduced (Fig. 1F,G). Conversely, the stabilizing E45A CA mutation reduced cGAS activation, resulting in limited cGAMP production even at very low IP6 levels. Both stabilized and destabilized capsid mutants were less infectious than wt HIV-1 when pseudotyped reporter viruses were compared in THP-1 monocytic cells (S1 Fig. G). Our in vitro results indicate that capsid stability modulates cGAS recognition, with stable capsids protecting viral DNA from cGAS recognition better than unstable capsids.

### Capsid inhibitors trigger viral core disruption and cGAS activation in vitro

Given that capsid stability regulates innate immune detection of HIV-1, we hypothesized that we could exploit chemical disruption of the capsid to unmask cGAS sensing of reverse transcripts. Consistent with this idea, the small molecule capsid inhibitor, PF74, can increase innate immune sensing of HIV-1 in cell lines, although this inhibitor has relatively low affinity for the capsid and only blocks viral replication at micromolar concentrations^15,44^. Advances in drug development have led to lenacapavir/SUNLENCA (LEN), an FDA-approved capsid inhibitor that is 1000-fold more potent than PF74^45,46^. LEN binding reduces core particle elasticity^47^ and stabilizes hexamer contacts^48,49^, thereby flattening and rigidifying patches of the conical CA lattice and inducing ruptures^3,50^. We therefore tested whether LEN could “expose” viral reverse transcripts for detection in our vitro system.

Time course experiments to track production of reverse transcripts relative to naked plasmid DNA again showed robust recombinant cGAS activity for the plasmid control, but not ERT samples, despite comparable levels of plasmid and ERT DNA (Fig. 2A). However, addition of LEN dramatically increased cGAS activation in the virus samples at late time points when dsDNA accumulated (Fig. 2A, lower panel). After 16 h of ERT, LEN increased cGAS detection of HIV-1 DNA by more than an order of magnitude, and this effect was specific for the ERT products because LEN did not alter cGAS sensing of naked plasmid DNA.

**Figure 2.**
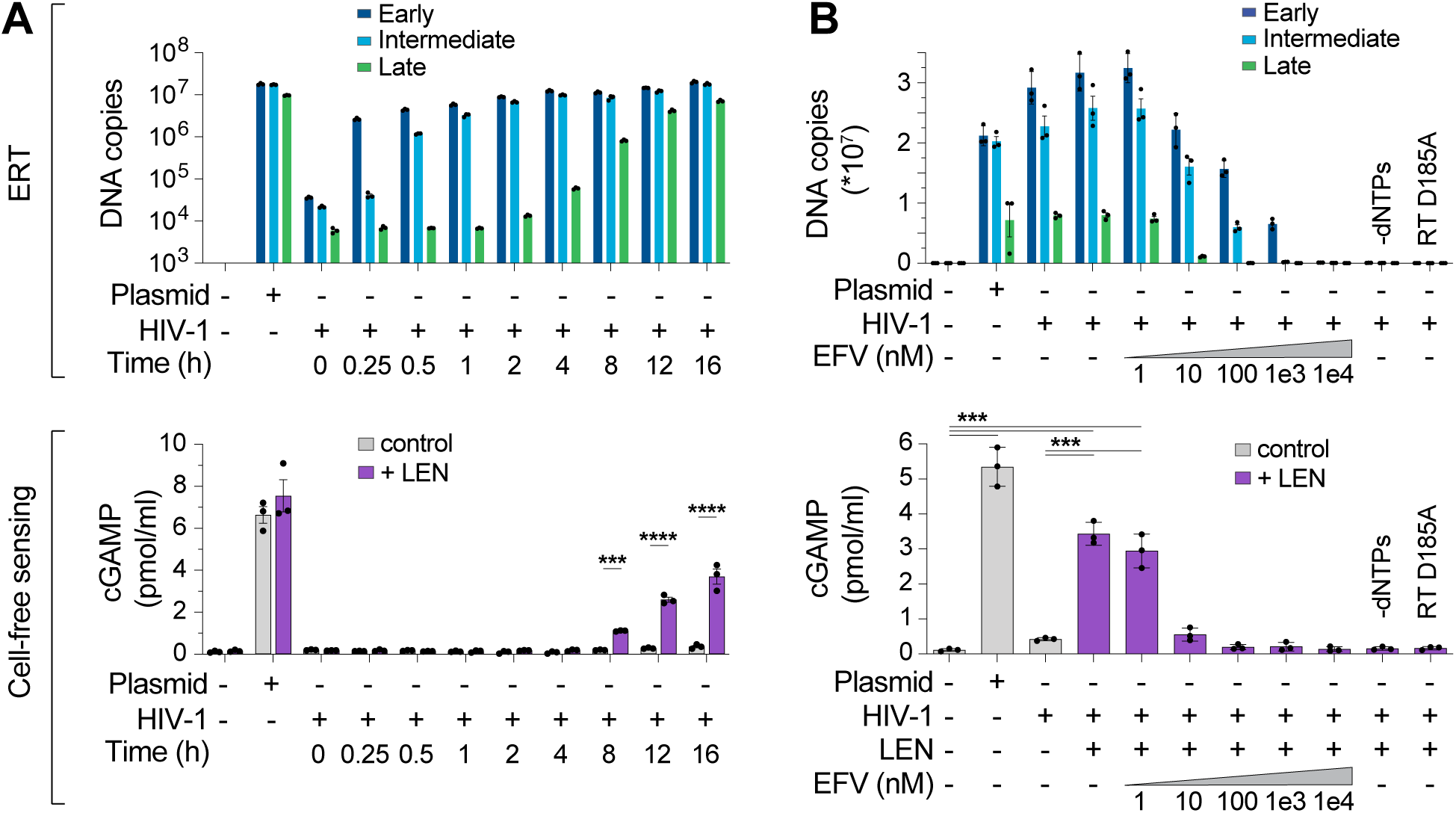
Lenacapavir triggers cGAS sensing of HIV-1 reverse transcripts. **(A)** DNA product levels from an HIV-1 ERT time course compared to a plasmid control, as measured by qPCR (top panel). After ERT, samples were incubated with recombinant cGAS ± LEN (100 nM) and cGAMP levels were determined by ELISA (cell-free sensing assays, bottom panel). **(B)** DNA products (top panel) from standard ERT reactions (16 h at 37°C) performed in the presence of increasing concentrations of the RT inhibitor efavirenz (EFV), or without dNTPs, or using a virus containing an inactivating mutation in RT (D185A). After ERT, samples were reacted with recombinant cGAS ± LEN (100 nM) and cGAMP levels were determined by ELISA (bottom panel). Statistics were calculated using a one-way ANOVA with Tukey’s multiple comparisons test: p<0.001: ***, p<0.0001: ****. Graphs depict mean ± SD from three samples from a representative experiment, selected from three independent experiments.

To determine whether cGAS activation by HIV-1 samples incubated with LEN depended on reverse transcription, we performed ERT and cGAS reactions with increasing concentrations of the reverse transcriptase inhibitor efavirenz (EFV). As expected, EFV blocked RT product formation in a dose-dependent manner (Fig. 2B, top panel). EFV treatments also reduced LEN-dependent cGAMP production, implying that cGAS activation reflected viral DNA sensing (Fig. 2B, bottom panel). Similar effects were observed for two other treatments that blocked ERT; omitting dNTPs or introducing a point mutation that inactivated reverse transcriptase (D185A) (Fig. 2B). Dropout experiments confirmed that, as expected, LEN-dependent cGAS sensing of HIV-1 RT products also required virion permeabilization (melittin), ERT (MgCl_2_ and IP6), and enzymatic cGAMP production (NTPs and cGAS) (S2 Fig. A-E). We note that higher concentrations of MgCl_2_ were associated with greater cGAS activity at baseline and reduced LEN responsiveness. We speculate that these MgCl_2_ effects could be driven by two factors: 1) cations such as Mg^2+^ and Mn^2+^ support cGAS activity in vitro^51^, and 2) at high concentrations, Mg^2+^ cations might disrupt capsid stability, by interfering with CA subunit interactions and/or by chelating IP6 away from the capsid, leading to loss of capsid integrity, genome release, and reduced LEN responsiveness. Importantly, cGAS sensing was also triggered by the LEN analog, GS-CA1^45^, and by the weaker PF74 inhibitor^52^, which all target the same capsid binding pocket^46,53^ (S2 Fig. F,G). Together, these data indicate that: 1) the HIV-1 capsid protects the viral genome from innate immune sensing, and 2) pharmacological disruption of the capsid enables cGAS detection of reverse transcripts.

We next assessed the specificity of the LEN effects by testing activity against CA mutations that inhibit LEN binding and confer drug resistance^46,54^. Control experiments confirmed that the M66I and Q67H/N74D CA mutations conferred LEN resistance to reporter virus infection of THP-1 cells, as expected (S3 Fig. A). In vitro, HIV-1 cores carrying these LEN resistance mutations completed ERT at normal levels (Fig. 3A). However, the mutant virions did not stimulate cGAS activity when treated with LEN (Fig. 3B). In control experiments, heat treated mutant capsids stimulated cGAS normally, indicating that their DNA was detectable when exposed by other methods of capsid disruption (S3 Fig. B). These experiments demonstrate that LEN binds the viral capsid to stimulate cGAS activity.

**Figure 3.**
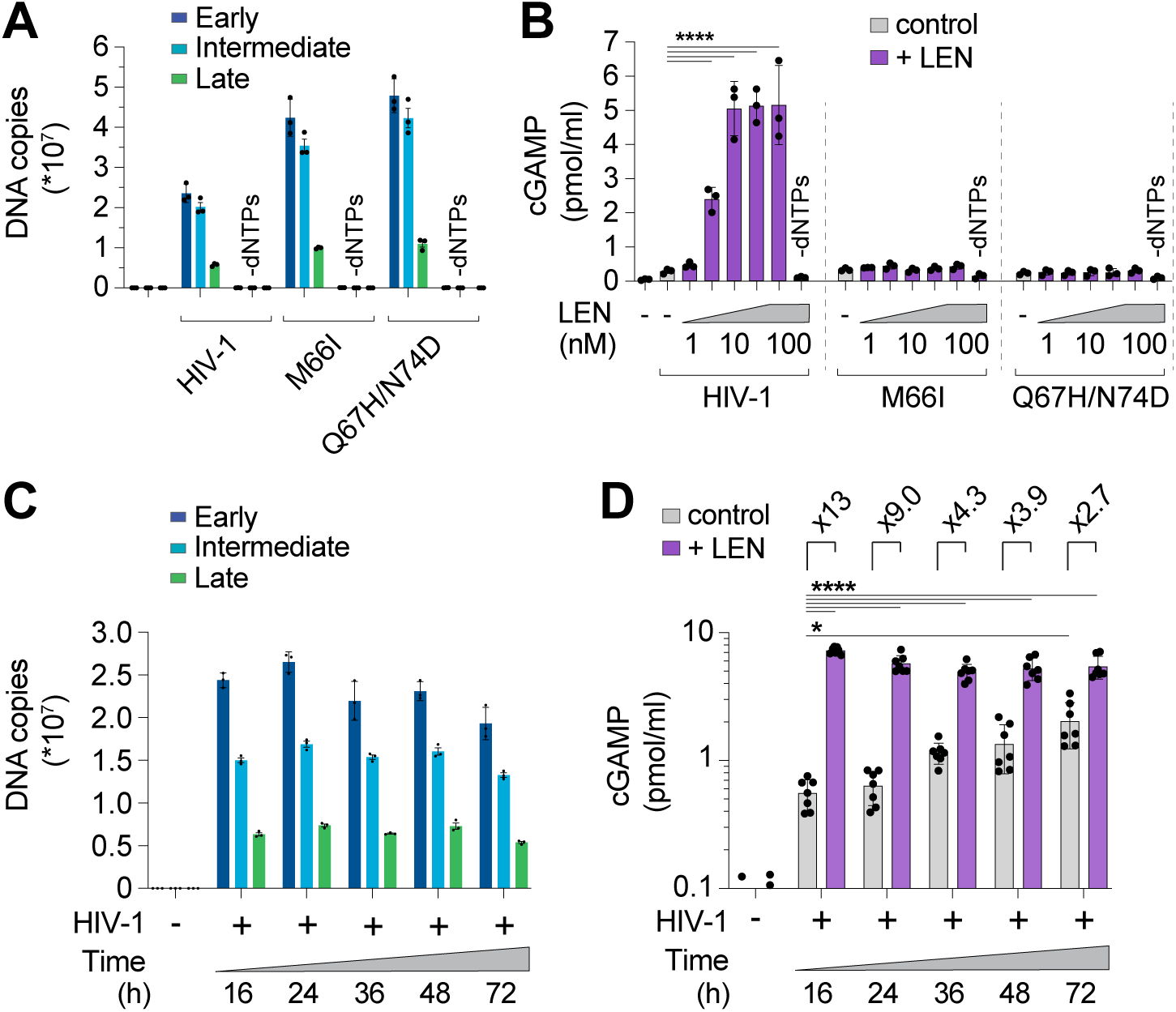
CA mutations confer resistance to LEN-triggered cGAS activation. **(A)** qPCR measurements of DNA copy number from ERT reactions under standard conditions (16 h at 37°C) with wt HIV-1 virions compared to virions bearing LEN resistance-associated mutations in CA (M66I and Q67H/N74D), performed with or without dNTPs. **(B)** Cell-free sensing assay of wt, M66I, or Q67H/N74D samples from (A). After ERT, samples were incubated with recombinant cGAS and increasing amounts of LEN (0, 1, 3.2, 10, 32, 100 nM), or LEN (100 nM) −dNTPs. cGAMP levels were determined by ELISA. **(C)** qPCR measurements of DNA copy number for ERT reactions that were held at 37°C for the indicated times. **(D)** Cell-free sensing assay of ERT samples from (C). After ERT, samples were normalized for late RT DNA copy number and then incubated with recombinant cGAS under standard conditions (± LEN at 100 nM). Brackets above graph represent the fold increase in cGAS activity with LEN at each time point. Statistics were calculated using a one-way ANOVA with Tukey’s multiple comparisons test: p<0.05: *, p<0.0001: ****. Graphs depict mean ± SD from three measurements from a representative experiment, selected from 3 independent experiments (A–C) or from 7 independent samples measured on different days (D).

During a normal infection, the viral DNA must eventually exit the capsid to integrate into the host genome, but the mechanism of release (termed “uncoating”) is not well understood. In particular, it is not yet clear whether uncoating occurs stochastically, is driven by pressure caused by accumulation of viral dsDNA, or is facilitated by host factors. To determine whether prolonged incubation of viral cores might lead to capsid permeability and genome release, we ran ERT reactions and incubated the samples for 16, 24, 36, 48, and 72 h at 37°C and then performed cGAS DNA sensing assays. DNA product levels were quite stable across these time points (Fig. 3C), and cGAS activation increased modestly (3-to 4-fold) from 16 to 72 h, indicating that ERT products became more exposed over time (grey bars, Fig. 3D). Nevertheless, even after 72 h, LEN treatment further stimulated cGAS activity (∼3-fold), indicating that capsids provide stable protection of viral DNA in vitro for days after completion of reverse transcription.

### LEN treatment triggers cellular sensing of HIV-1 via the cGAS-STING pathway

We also extended our findings to cellular models of HIV infection. In vitro, LEN-dependent cGAS activation by ERT reactions exhibited biphasic behavior (Fig. 4A). At high concentrations (>30 nM), LEN inhibited reverse transcription (Fig. 4A, top panel) and cGAS activation was therefore prevented because dsDNA was not present for detection^3^. Lower LEN concentrations (1–10 nM) stimulated cGAS activation in a dose dependent fashion, consistent with the idea that reverse transcription proceeded, and viral cores were increasingly disrupted, triggering cGAS activation (Fig. 4A, bottom panel). To determine whether these effects were recapitulated in the context of a viral infection, we tested the effects of different LEN levels on THP-1 cells treated with HIV-1-GFP and tracked both infectivity and innate immune activation, as measured by induction of two different interferon-stimulated genes (ISGs): ISG15 and SIGLEC1 (Fig. 4B,C). We found that high LEN levels inhibited cGAS activation (likely due to reduced reverse transcription, as described below), but lower LEN levels stimulated ISG15 and SIGLEC1 expression (Figs. 4C; S4 Fig. A-E), at concentrations that matched the levels that stimulated cGAS activity in vitro (1–10 nM). Thus, the LEN effects observed in cells mirror those observed in vitro.

**Figure 4.**
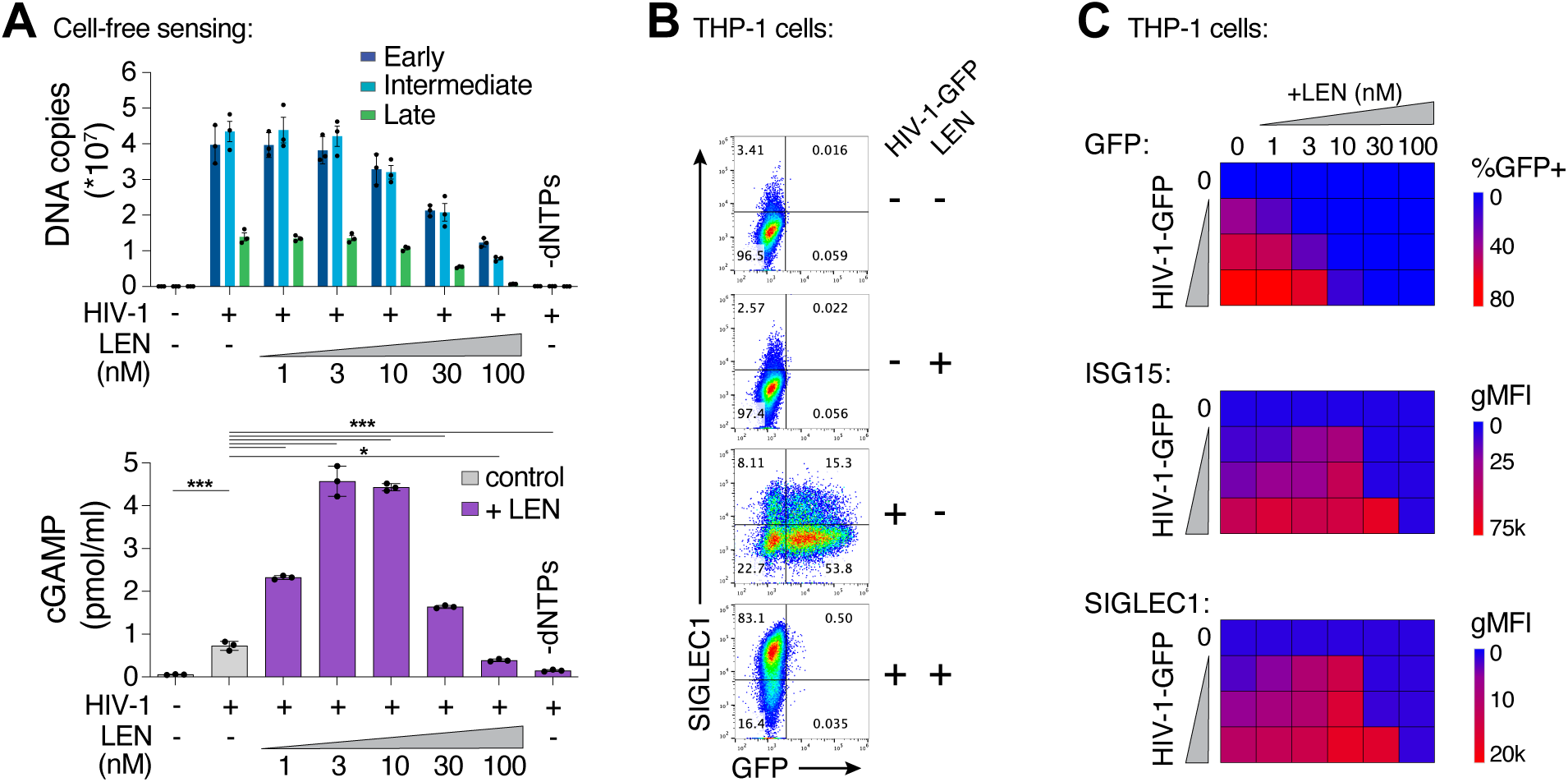
LEN potentiates sensing of HIV-1 in vitro and in cells. **(A)** qPCR measurements of DNA copy number from ERT reactions performed under standard conditions (16 h at 37°C) with or without LEN at the indicated concentrations (top panel). After ERT, samples were reacted with recombinant cGAS in cell-free sensing assays (bottom panel), maintaining the same concentrations of LEN as used in ERT. **(B)** Flow cytometry of THP-1 monocytic cells infected with HIV-1-GFP (12.5 nM p24) for 48 h (± LEN at 10 nM), showing expression of GFP (viral infectivity) and SIGLEC1 (IFN response). **(C)** Heat maps of GFP, ISG15, and SIGLEC1 expression as determined by flow cytometry of THP-1 cells infected with HIV-1-GFP for 48 h. Columns show a range of LEN concentrations and rows show increasing virus doses (12.5, 25, and 50 nM p24). n = 3. For (A), statistics were calculated using a one-way ANOVA with Tukey’s multiple comparisons test: p<0.05: *, p<0.001: ***, p<0.0001: ****. Graphs depict mean ± SD from three samples from a representative experiment, selected from 2–3 independent experiments, unless otherwise noted.

We confirmed that high concentrations of LEN (>30 nM) reduced early, intermediate, and late HIV-1-GFP reverse transcripts in THP-1 cells (Fig. 5A), which agrees with recently published work^33^. Our dose response matrix of virus and LEN concentrations indicated that potent innate immune responses could be revealed with low virus input and therapeutic levels of LEN (e.g. 10 nM, S4 Fig. A-B). To validate these findings further and evaluate the kinetics of the innate response, we tested HIV-1-GFP at low mutliplicity of infection (MOI) and measured ISG expression at different times after infection. THP-1 cells began to express GFP roughly 24 h after exposure to low levels of HIV-1-GFP (12.5 nM p24; MOI < ∼0.5) but did not express high levels of ISGs during the two-day time period tested (Fig. 5B). In contrast, the same virus dose in the presence of 10 nM LEN triggered strong induction of a series of ISGs (*IFITM1, IFITM3, MX1, ISG15*, and *IFI27*), indicating that LEN exposed viral DNA above the threshold required for cGAS-STING activation.

**Figure 5.**
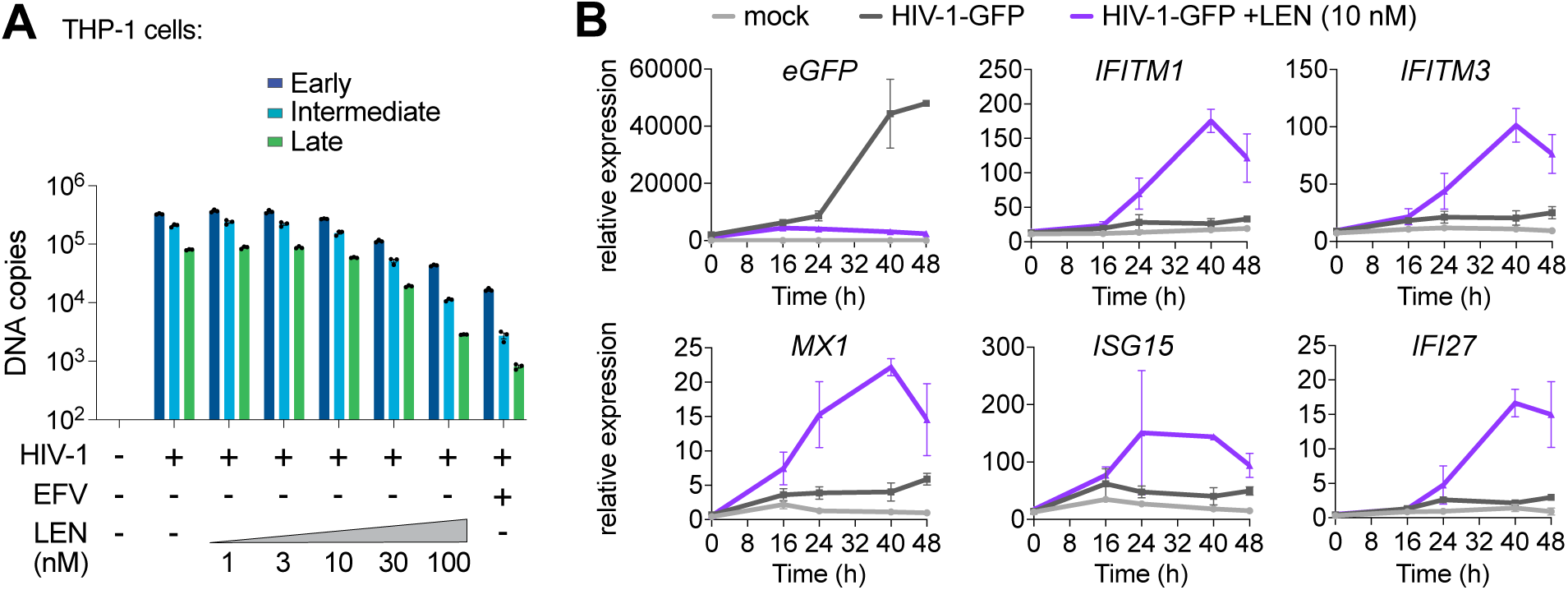
Therapeutic concentrations of LEN activate innate immune sensing at low MOI. **(A)** qPCR measurements of reverse transcription in THP-1 monocytic cells that were infected for 24 h with HIV-1-GFP (12.5 nM p24, corresponding to an MOI of < 0.5) with increasing concentrations of LEN. EFV (10 µM) was included as a positive control to inhibit RT. **(B)** Time course of gene expression in THP-1 monocytic cells infected with HIV-1-GFP (12.5nM p24) with or without LEN (10 nM). Graphs show qPCR measurements of relative expression for *eGFP, IFITM1, IFITM3, MX1, ISG15*, and *IFI27* at 0, 16, 24, 40, and 48 h time points. Graphs depict mean ± SD from three samples from a representative experiment, selected from 3 independent experiments.

LEN treatments did not stimulate ISG expression upon infection by the LEN-resistant mutants M66I and Q67H/N74D, showing the specificity of drug activity in cells (S4 Fig. F). We note, however, that although the M66I mutation reduced viral infectivity, the mutant virus induced ISG expression at levels that were similar to wt HIV-1, consistent with the idea that this mutation reduces viral fitness owing to capsid structural defects^49,55^. Most importantly, our data demonstrate that LEN can stimulate innate immune sensing of incoming wt HIV-1 particles, at therapeutically relevant concentrations that still block infection.

To determine whether LEN-triggered innate immune activation occurred through the canonical cGAS/STING pathway, we employed the lentiCRISPR system^56^ to knock out the key pathway components, cGAS, STING, or IRF3. Pools of THP-1 monocytic cells were transduced with lentiCRISPR vectors, expanded for two weeks under antibiotic selection, and target protein loss was confirmed by immunoblot (Fig. 6A). As expected, HIV-1 infection was blocked by LEN treatment (10 nM), but was not affected by loss of cGAS, STING, or IRF3 (Fig. 6B). However, loss of each pathway component substantially reduced ISG protein expression, both in the absence and presence of LEN (Fig. 6C). Thus, LEN stimulates innate immune recognition of HIV-1 through the canonical cGAS/STING pathway.

**Figure 6.**
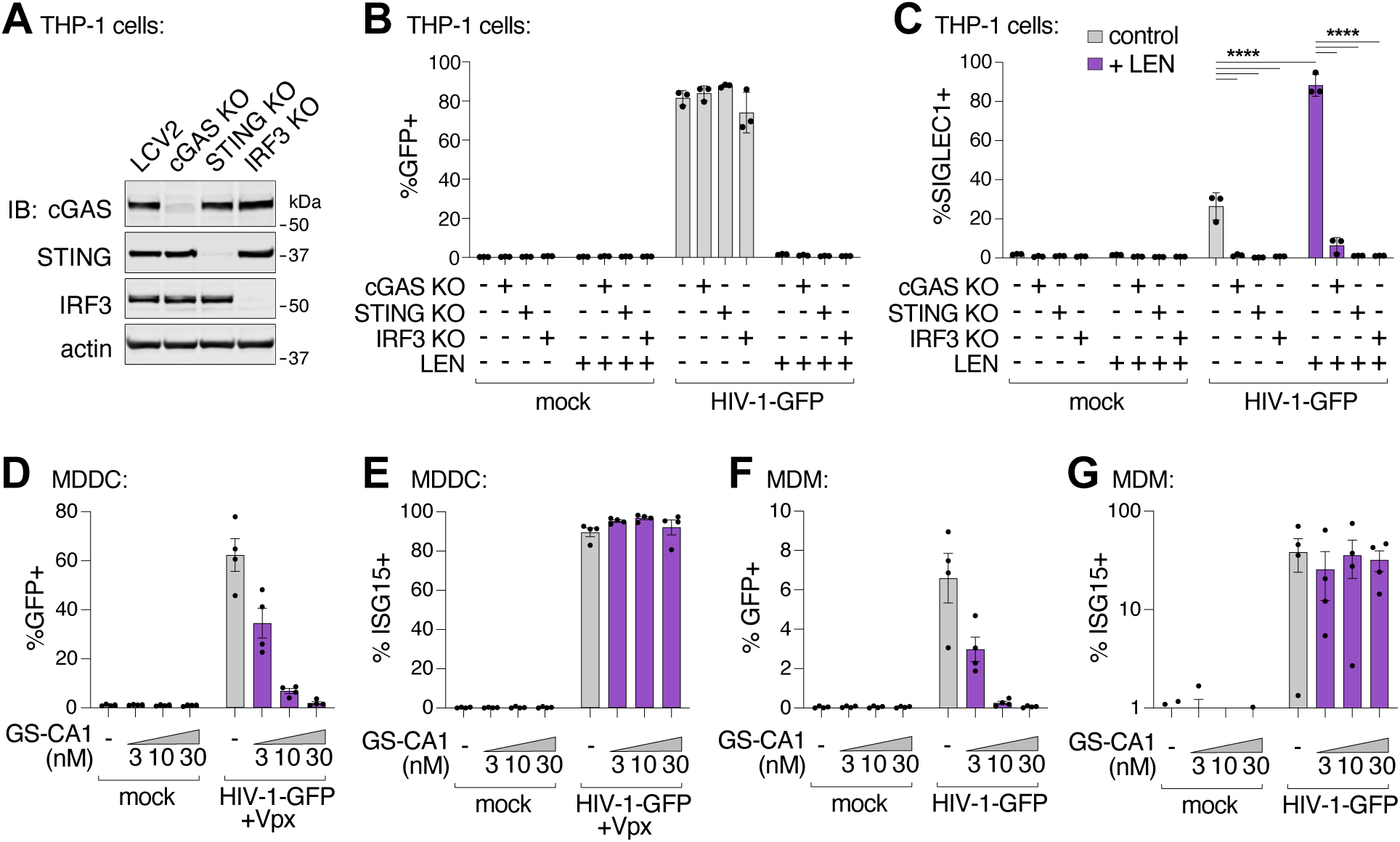
Capsid inhibitors potentiate sensing of HIV-1 through the cGAS-STING pathway in cells. **(A)** Immunoblots of THP-1 lysates after lentiCRISPR editing, depicting knockout of cGAS, STING, or IRF3 relative to a non-targeting vector control (LCV2). **(B–C)** Flow cytometry of lentiCRISPR-modified THP-1 cells infected with HIV-1-GFP (25 nM p24) for 48 h, showing the percent of cells positive for GFP (B) or SIGLEC1 (C) expression for the indicated conditions (± LEN at 10 nM). **(D–E)** Flow cytometry of MDDCs (derived from 4 separate donors) that were infected with HIV-1-GFP for 48 h with increasing concentrations of GS-CA1 (0, 3, 10, and 30 nM) added at the time of infection. Vpx was included to overcome SAMHD1-mediated restriction. Panels show infectivity (GFP+, D) and IFN induction (ISG15+, E). **(F–G)** Flow cytometry of MDMs (derived from 4 donors) that were infected with HIV-1-GFP in the same fashion as in (D–E) with increasing concentrations of GS-CA1 (0, 3, 10, and 30 nM) added at the time of infection. Infections were performed without Vpx. Graphs depict the percentage of GFP (D,F) or ISG15 (E,G) positive cells. For (C), statistics were calculated using a one-way ANOVA with Tukey’s multiple comparisons test: p<0.05: *, p<0.001: ***, p<0.0001: ****. Graphs depict mean ± SD from three samples from a representative experiment, selected from 2–3 independent experiments, unless otherwise noted.

Recent work has suggested that an additional factor, polyglutamine-binding protein 1 (PQBP1), helps to license cGAS activity during HIV-1 infection, through its capacity to link cGAS to the viral capsid^32,57^. In our cell-free system, full-length recombinant PQBP1 protein did not enhance cGAS activity, either alone or together with HIV-1 (S5 Fig. A,B). Similarly, we observed that loss of PQBP1 did not alter viral infectivity or reduce innate immune sensing of HIV-1 in myeloid cells, using two independent guide RNAs to knockout PQBP1 expression (S5 Fig. C-G). Loss of PQBP1 also had no effect when innate immune sensing was limited to early steps of the life cycle in the presence of the integration inhibitor elvitegravir (S5 Fig. F,G). In contrast, cGAS removal (positive control) ablated ISG responses to HIV-1-GFP under all conditions tested.

Finally, we evaluated whether capsid inhibition could facilitate innate immune responses while blocking HIV-1 infection in primary human myeloid cells. Human monocyte-derived dendritic cells (MDDCs) and monocyte-derived macrophages (MDMs) can be infected by HIV-1, particularly when the lentiviral accessory protein Vpx is provided in trans to overcome SAMHD1-mediated restriction in MDDCs^58,59^. In some cases, MDDCs and MDMs can respond to HIV-1 by producing type I IFN and other inflammatory cytokines^16,17,19,21,60,61^. We infected MDDCs and MDMs from four unique donors with HIV-1-GFP (Fig. 6D,F), and found that 3–30 nM concentrations of the LEN analog, GS-CA1, efficiently blocked infection in MDDCs and MDMs. Importantly, however, ISG15 activation was still sustained at these inhibitory concentrations (Fig. 6E,G). Hence, potent LEN-class capsid inhibitors can simultaneously prevent HIV-1 infection while still maintaining robust innate immune responses in primary human myeloid cells.

## Discussion

IFN creates strong selective pressure on HIV-1 during sexual transmission and during viral rebound in response to withdrawal of antiviral therapy^62,63^. It is therefore critical to understand how HIV is sensed by the innate immune system. Multiple innate immune sensors can contribute to IFN and inflammation during HIV-1 infection^60,61,64–68^, but cGAS activation is increasingly appreciated as a driver of protective anti-HIV immune responses. For example, this pathway is robustly activated in myeloid cells from elite controllers^69,70^, and myeloid cell programming of polyfunctional CD8+ T cells is critical for control of infection^71,72^. However, precisely how viral DNA is released from the capsid to activate cGAS has not been well characterized.

To begin to address this issue, we have reconstituted endogenous reverse transcription, capsid destabilization, and innate immune sensing of HIV-1 in a cell-free system. We find that HIV-1 capsids protect reverse transcripts from detection by cGAS, and that this protection can last for days after reverse transcription is complete. This protection appears to be mediated by intact, or largely intact capsids because cGAS recognition increases dramatically when capsids are disrupted by heat, reduced IP6 concentrations, or destabilizing mutations in the viral capsid. Our observation that capsid integrity and stability can strongly modulate cGAS-mediated detection of HIV-1 is consistent with a growing body of literature indicating that the capsid sequence has a substantial impact on sensitivity to innate immune sensors^16,17,21,28,30^, and that IP6 perturbations or improper capsid formation can promote cGAS activation^15,29^. We suggest that our in vitro system can be useful in characterizing these effects, and also in determining how host factors promote capsid uncoating and cDNA exposure.

Our work demonstrates that wt HIV-1 capsids have the intrinsic property of shielding viral DNA from innate immune detection, but we note that recruitment of host factors likely serves to further modify capsid stability in cells (reviewed in ^73,74^). For example, cyclophilin A (CypA)^75^ and CPSF6^76^ are well-characterized CA-binding proteins that play different roles during entry and may positively or negatively affect the capsid, as they can promote trafficking^77^, influence nuclear entry^78,79^, and affect integration site selection^80,81^. Another protein, TRIM5α from rhesus macaques can form a hexagonal cage surrounding the HIV-1 capsid^82^ and block early steps in infection^83,84^, potentially by disrupting core integrity or other mechanisms^85^.

Moving forward, our cell-free assay will likely be useful for determining whether CypA, CPSF6, TRIM5α, and other capsid-binding factors have a direct effect on core stability. In this study, we tested the role of PQBP1, as it has been reported to bind to both the capsid and to cGAS, and to be important for innate immune sensing of HIV-1^32,57^. Although we did not observe an effect with PQBP1 in vitro or validate that it is required for HIV-1 detection in myeloid cells, we cannot rule out that PQBP1 plays an important role under different circumstances.

Importantly, we found that capsid inhibitors of the LEN class can promote HIV-1 DNA genome exposure and cGAS activation in vitro, in THP-1 cells, and in primary human myeloid cells. In the THP-1 case, and presumably also in primary myeloid cells, cGAS activation by reverse transcription products leads to innate immune activation via the canonical STING pathway (Fig. 6), which supports the growing body of literature evoking the importance of this pathway in HIV innate immunity^15–19,28–31,33,57,69,70^. However, we have not yet identified conditions that lead to LEN-induced IFN production in CD4+ T cells, suggesting that additional host circuitry might be required to unleash this response in the main target cells of HIV^86,87^.

In summary, our experiments are consistent with a model in which viral cores enter the cytosol of myeloid cells, and reverse transcription normally proceeds inside intact or largely intact capsids, where viral cDNA is protected from innate immune sensors^1,16,21,31^. However, in the presence of LEN, accumulating reverse transcripts can become exposed by drug-induced capsid rupture, enabling viral DNA detection and downstream activation of IFN production via the cGAS-STING-IRF3 pathway (S1 Movie). Importantly, LEN can inhibit multiple steps in the virus life cycle at different drug concentrations, including virus assembly and a step corresponding to nuclear entry or a later pre-integration step, with the latter being the most potent block to infection in cultured cells (0.5–5 nM)^45^. Reverse transcription can also be inhibited, but this requires drug concentrations (≥25 nM) that are much higher than the protein-adjusted EC95 required to inhibit HIV-1 replication (∼4 nM)^45,46^. We found that LEN promotes cGAS activation at concentrations that parallel those that block nuclear entry and/or integration (1–10 nM, see Fig. 4), and we hypothesize that the most potent mechanism of drug action may be associated with premature genome exposure. Our experiments also indicate that, at least in culture, cGAS activation can occur under conditions where viral infectivity is suppressed but innate immune responses are maintained. We therefore speculate that the drug-dependent effects we observe in cultured cells might be harnessed to potentiate vaccination responses under conditions where viral replication is blocked while antiviral immunity remains engaged.

## Acknowledgments

We thank members of the Johnson and Sundquist laboratories, Owen Pornillos, Barbie Ganser-Pornillos, Jody Puglisi, Elisabetta Viani Puglisi, and other CHEETAH Center investigators for helpful discussions. Oligonucleotides were synthesized by the University of Utah DNA/Peptide Core Facility, sequencing of constructs was performed at the University of Utah DNA sequencing Core Facility, and flow cytometry was performed at the University of Utah Flow Core. We acknowledge funding from National Institutes of Health grant U54 AI170856 (to WIS and JSJ), R56 AI174930 (JSJ), R01 AI181713 (JSJ), 1DP2GM146250-01 (PJK), and grants from the Pew Biomedical Scholars program (PJK), and the Burroughs Wellcome Fund PATH program (PJK). W.Z. was supported in part through a Charles A. King Trust Postdoctoral Fellowship. The funders had no role in study design, data collection, and interpretation, or the decision to submit the work for publication.

## Author contributions

Conceptualization: WIS, JSJ

Methodology: TMS, LMA, DEC, MJP, WZ, PJK, WIS, JSJ

Investigation: TMS, LMA, JAP, DAM, ABR, DEC, WZ, JSJ

Funding acquisition: PJK, WIS, JSJ

Project administration: PJK, WIS, JSJ

Supervision: PJK, WIS, JSJ

Writing – original draft: JSJ

Writing – review & editing: all authors

## Competing interests

The authors declare no competing interests.

## Methods

### Cell lines and blood-derived dendritic cells

To generate MDDCs and MDMs, we acquired leukocytes from de-identified normal human donors from ARUP Blood Services, Sandy, UT, USA, similar to previously described methods^88^. We cannot report on the sex, gender, or age of the donors since the samples were de-identified and donors remain anonymous. Peripheral blood mononuclear cells (PBMCs) were layered over Ficoll-Paque Plus (GE Healthcare). CD14+ monocytes from PBMC buffy coats were isolated with anti-human CD14 magnetic beads (Miltenyi) and cultured in RPMI containing 10% heat-inactivated fetal bovine serum (FBS, Gibco), 50 U/mL penicillin, 50 μg/mL streptomycin (pen/strep, Thermo Fisher), 10 mM HEPES (Sigma), and 55 μM 2-Mercaptoethanol (Gibco). We tested multiple lots of FBS to identify batches that lead to minimal baseline induction of activation markers over the course of MDDC differentiation. MDDCs were derived in the presence of recombinant human GM-CSF (Peprotech) at 10 ng/mL and IL-4 at 50 ng/mL. MDMs were derived using recombinant human M-CSF (Peprotech) at 20 ng/mL. Fresh media and cytokines were added to cells (40% by volume) one day after CD14+ cell isolation. On day 4, MDDCs were collected, resuspended in fresh media with cytokines used for infection or stimulation such that experimental end points would fall on day 6. MDM cultures were replenished with fresh media on day 4, then infected or stimulated on day 5 so that experimental end points would fall on day 7 after isolation. MDDC and MDM experiments were performed using biological replicates from blood-derived cells from multiple individual donors as indicated in the figure legends.

We cultured 293FT cells (immortalized from a female fetus, Life Technologies Cat# R70007, RRID: CVCL_6911) in Dulbecco’s modified Eagle’s medium (DMEM, Thermo Fisher) supplemented with 10% heat-inactivated FBS, pen/strep, 10mM HEPES, and 0.1 mM MEM non-essential amino acids (Thermo Fisher). THP-1 cells (derived from a male patient with acute monocytic leukemia, ATCC) were cultured in RPMI (Thermo Fisher) containing 10% FBS, pen/strep, 10 mM HEPES, and 2-Mercaptoethanol.

Cell lines were used at early passage numbers and fresh stocks were thawed every 1–2 months. Cell cultures were routinely tested for mycoplasma contamination every 6 months. All cells were maintained at 37°C and 5% CO_2_.

### Plasmids and mutagenesis

For endogenous reverse transcription reactions, HIV-1_SG3_ was produced from pSG3Δenv plasmid, which was acquired from the NIH AIDS Reagent Program (Catalog # 11051) and contains mutations in the *env* gene rendering it non-functional (ΔEnv), but otherwise represents the full-length HIV-1 genome. HIV-1-GFP is derived from an HIV-1 NL4-3 vector that has been used to study immune responses in myeloid cells lines and primary human MDDCs^21^. pHIV-1-GFP is *env-vpu-vpr-vif-nef-*, with the GFP open reading frame in place of *nef*. We generated virus like particles packaging Vpx from the plasmid pSIV3+ (based on SIVmac251, GenBank acc. no. M19499), which has been described previously^89^. These and all other plasmids used in this study are listed in the Supplementary Table (S1 Table) and have been made available through Addgene. Target sequences for lentiCRISPR vectors are also provided (S1 Table). All lentiviral constructs were transformed into Stbl3 bacteria (Thermo Fisher) for propagation of plasmid DNA. The coding sequence for full-length human PQBP1 was synthesized as a GBlock gene fragment (IDT) and cloned into pCA528 by Gibson Assembly. All plasmids were prepped and purified through PureLink HiPure Plasmid Maxiprep Kits (Invitrogen).

### Virus and vector production

Reporter viruses, recombinant lentiviral vectors, and Vpx-containing virus-like particles, were produced by transient transfection in 293FT cells, similar to previously described methods ^19^. Briefly, the day before transfection, cells were seeded onto poly-L-lysine-(Sigma) coated 15 cm plates to be ∼70% confluent at the time of transfection. Cells were transfected with a total of 22–25 μg DNA using PEImax (Polysciences, Inc.). Each batch of PEI was titered to identify the optimal ratio of DNA:PEI for transfection (which was typically close to 1:2). To produce wt SG3ΔEnv and mutant virions, a single plasmid was transfected into 293FT cells. For HIV-1-GFP reporter viruses, we transfected 19.1 μg of the HIV cassette and 3.4 μg of pCMV-VSV-G. For lentiCRISPR vectors, we transfected 13.5 μg of the transgene plasmid, 8 μg of psPax2 helper plasmid, and 3.5 μg pCMV-VSV-G. Virus-like particles containing Vpx were produced by transfecting 19.1 μg pSIV3+ and 3.5 μg pCMV-VSV-G. The morning after transfection, cells were washed once and replenished with 30 mL of fresh media. Supernatants were harvested 36–48 h after replenishing media and then passed through 0.45 μm syringe filters (Corning) to remove debris.

To purify wt and mutant SG3ΔEnv virions used for ERT reactions and cell-free sensing assays, virus supernatants (30 mL) were treated for 1 h at 37°C with 2500 Units ultrapure benzonase nuclease (Sigma, E8263-25KU) and subtilisin^90^ (Sigma). Protease activity was then quenched with PMSF. Virus preparations were further purified through methods that will be described in a forthcoming publication. HIV-1-GFP wt and mutant virions were treated in a similar fashion except were not treated with subtilisin and PMSF. Supernatants were layered over a 20% sucrose cushion (in PBS) and concentrated by ultracentrifugation by spinning in 25 x 89 mm ultraclear tubes (Beckman) at 28 K rpm for 2 h at 4°C in an SW32 swing-bucket rotor (Beckman). SG3ΔEnv virions were thoroughly resuspended using 500 μL Hepes/Salt buffer per tube. HIV-1-GFP wt and mutant virions and lentiCRISPR vectors were resuspended in 1 mL of complete RPMI media. With high titer stocks of HIV-1-GFP and lentiCRISPR vectors we occasionally observed insoluble virus aggregates that we clarified from virion preparations by low-speed centrifugation at 300 rcf for 3 min at 4°C. Purified viral stocks were frozen at −80°C, titered by p24 ELISA (Express Bio), and used as indicated below.

### Bacterial expression and purification of recombinant human cGAS

Full-length human cGAS (aa 1–522, HGNC:21367) was produced and purified as previously described^91,92^. Briefly, human cGAS was expressed as a fusion protein with a 6×His-SUMO2 tag in *Escherichia coli* (*E. coli*) BL21-RIL DE3 (Agilent) bacteria harboring a pRARE2 tRNA plasmid. Initial cultures were grown in 30 mL of MDG media (25 mM Na_2_HPO_4_, 25 mM KH_2_PO_4_, 50 mM NH_4_Cl, 5 mM Na_2_SO_4_, 2 mM MgSO_4_, 0.5% glucose, 0.25% aspartic acid, 100 μg/mL ampicillin, 34 μg/mL chloramphenicol, and trace metals) to an OD_600_ of ∼0.05. These cultures were then used to seed 2 L M9ZB media (7.8 mM Na_2_HPO_4_, 22 mM KH_2_PO_4_, 18.7 mM NH_4_Cl, 85.6 mM NaCl, 2 mM MgSO_4_, 0.5% glycerol, 1% Casamino acids, 100 μg/mL ampicillin, 34 μg/mL chloramphenicol, and trace metals) at 37°C until an OD_600_ of ∼2.5–3.0 was reached. The cultures were supplemented with Isopropyl ß-D-1-thiogalactopyranoside (IPTG) (0.5 mM) and incubated at 16°C for an additional 12–16 h. Next, bacteria were pelleted, flash-frozen in liquid nitrogen, and stored at −80°C until further purification.

Bacterial pellets with cGAS fusion proteins were resuspended in lysis buffer (20 mM HEPES-KOH pH 7.5, 400 mM NaCl, 10% glycerol, 30 mM imidazole, 1 mM dithiothreitol (DTT) and subsequently lysed by sonication. The supernatant containing cGAS was collected for initial purification using Ni-NTA (Qiagen) resin and gravity chromatography. Eluted 6×His-SUMO2-cGAS proteins were supplemented with ∼250 μg of human SENP2 protease to remove the His-SUMO2 tag and treated in dialysis buffer (20 mM HEPES-KOH pH 7.5, 300 mM NaCl, 1 mM DTT) at 4°C for ∼12 h. The dialysate was purified by using heparin HP ion exchange (GE Healthcare) and eluted with a 300–1000 mM NaCl gradient. Full-length untagged cGAS protein was then ultimately purified using size-exclusion chromatography via a 16/600 Superdex S75 column (GE Healthcare). Purified cGAS was concentrated to ∼10 mg/mL in storage buffer (20 mM HEPES-KOH pH 7.5, 250 mM KCl, 1 mM TCEP), flash-frozen in liquid nitrogen, and stored as aliquots at −80°C for biochemical experiments.

### Bacterial expression and purification of recombinant human PQBP1

Full-length human PQBP1 (HGNC:9330) was cloned into pCA528 for bacterial expression as a (His)_6_-SUMO-fusion protein. (His)_6_-SUMO-PQBP1 was expressed in BL21 RIPL cells grown in 2xYT media. Transformed cells were initially grown for 3–6 h at 37°C to an OD600 of 0-4–0.6, IPTG was added to final concentration of 0.5 mM to induce recombinant protein expression and cells were cultured for an additional 4 h. Cells were harvested by centrifugation at 6,000 x g and cell pellets were stored at −80°C.

All purification steps were carried out at 4°C except where noted. Frozen cell pellets were thawed and resuspended in lysis buffer: 50 mM Tris (pH 8.0 at 23 °C), 500 mM NaCl, 2 mM Imidazole, 0.5 mM EDTA, supplemented with lysozyme (25 µg/mL), PMSF (100 µM), Pepstatin (10 µM), Leupeptin (100 µM), Aprotinin (1 µM), and DNase I (10 µg/mL). Cells were lysed by sonication and lysates were clarified by centrifugation at 40,000 x g for 45 min. Clarified supernatant was filtered through a 0.45 µM PES syringe filter and incubated with 10 mL of cOmplete His-Tag purification beads (Roche) for 1 h with gentle rocking. Beads were washed with 150 mL unsupplemented lysis buffer. PQBP1 fusion protein was eluted with 50 mL of lysis buffer supplemented with 250 mM Imidazole.

Eluted proteins were treated with 100 μg His_6_-ULP1 protease overnight in 6–8 k MWCO dialysis bags while dialyzing against 2 x 2 L of 25 mM Tris (pH 8.0 at 23°C), 50 mM NaCl, 1 mM TCEP. The dialysate was purified by Q Sepharose chromatography (HiTrap Q HP 5 mL; Cytiva Life Sciences) with a linear gradient elution from 50–1000 mM NaCl. Fractions containing processed fusion proteins were then passed over 5 mL of cOmplete His-Tag purification beads to remove residual uncleaved His_6_-Sumo-PQBP1 and His_6_-Sumo cleaved tag. The sample was then placed in 6–8 k MWCO dialysis bag while dialyzing against 2 x 2 L of 25 mM Tris pH (8.0 at 23°C), 50 mM NaCl, 1 mM TCEP. The dialysate was purified by Heparin Sepharose chromatography (HiTrap Heparin HP 5 mL; Cytiva Life Sciences) with a linear gradient elution from 50–1000 mM NaCl. PQBP1 rich fractions were concentrated and purified by Superdex 75 gel filtration chromatography (120 mL;16/600; Cytiva Life Sciences) in 50 mM Tris (pH 8.0 at 23°C) 150 mM NaCl, 1 mM TCEP. Highly pure PQBP1 fractions were pooled, concentrated to ∼200 μM, aliquoted and stored at −80°C.

### Endogenous Reverse Transcription (ERT)

Experimental conditions for ERT reactions were optimized from existing protocols^3,36^.

Briefly, we diluted HIV-1_SG3Δ_Env virions to 50 nM CA in reaction mixtures that contained 50 mM Tris (pH 7.5 at 37°C), 100 mM NaCl, 1.65 mM MgCl_2_, 40 μM IP6, 1mg/mL bovine serum albumin (BSA), 10 μg/mL melittin (Sigma), ribonucleoside triphosphates (NTPs) (Promega), and deoxynucleotide triphosphates (dNTPs) (Promega). NTP concentrations were chosen to match levels reported in macrophage cytoplasm^93^ and were as follows: ATP: 1.124 mM; GTP: 323 μM; UTP: 173 μM; CTP: 25 μM. The concentration of Mg2+ was chosen to match the total concentration of NTPs in the reaction (1.65 mM) and we validated empirically that this optimized ERT yield. dNTP concentrations were previously established^3,94^ and were as follows: dATP: 5.2 μM; dGTP: 4.6 μM; dCTP: 6 μM; dTTP: 8 μM. We determined that ERT reactions could proceed using a range of melittin concentrations, and we chose 10 μg/mL for our standard conditions since higher concentrations of melittin (>25 μg/mL) were found to inhibit cGAS activity in the second stage of the assay. Reducing the concentration of melittin below 2.5 μg/mL led to reduced IP6 dependence (i.e. late RT products were produced even in the absence of IP6), suggesting that at lower concentrations of melittin, the membrane was incompletely stripped from the viral core and capsid stability was no longer influenced by IP6.

For plasmid comparisons in ERT reactions, we used the pSG3Δenv plasmid diluted to match the expected copy number for late reverse transcription products from HIV-1_SG3_ (typically 20–40 ng/reaction). Plasmid and HIV-1_SG3_ virions were incubated under identical conditions as described in the figure legends. After 16 h ERT incubations, column-based purification, and qPCR determination of DNA copy number from eluates (described below), plasmid copy numbers ranged from 5 M to 10 M copies per qPCR reaction. ERT reactions were performed in 60 μL volumes and held at 37°C in a ProFlex thermocycler (Thermo Fisher) for the times indicated in the figure legends and cooled to room temperature prior to downstream studies. Standard ERT reactions were performed for 16 h at 37°C. Samples from the same ERT reactions were divided into cGAS sensing assays or used for qPCR analysis of DNA yield.

To compare plasmid control DNA in circular and linear forms, a restriction digest of the pSG3Δenv plasmid was performed (± EcoRI-HF) using rCutSmart buffer (New England Biolabs). Samples were digested for 20 min at 37°C. Undigested and digested plasmid samples were separated on a 0.7% agarose gel for 1 h at 120V before extracting bands corresponding to circular/supercoiled and linearized DNA using a QIAquick Gel Extraction Kit (Qiagen). Circular and linear products were then added to ERT reactions as outlined above.

### Cell-free cGAS sensing assay

Samples from ERT reactions were diluted 50-to 100-fold into cell-free cGAS assay buffers that were similar to standard ERT reaction mixtures, but lacked melittin (50 mM Tris pH 7.5, 100 mM NaCl, 40 μM IP6, 1mg/mL BSA, NTPs, and 2.2 mM MgCl_2_). Full length recombinant human cGAS was added at 10 nM. cGAS assays were performed for 8 h at 37°C in a thermocycler, with reaction volumes typically ranging from 100-200 μL. We confirmed a linear response in signal for shorter and longer incubations with cGAS (4, 8, and 16 h). BSA was not required for DNA product formation in ERT reactions but was critical to observe a cGAMP signal above the limit of detection in cell-free cGAS assays. Product formation in the cGAS assay was exquisitely sensitive to Mg^2+^ concentration (S1 Fig. E). Although we observed similar responses to HIV-1 + LEN over a range of Mg^2+^concentrations (1.65 mM to 5 mM), our standard reaction conditions reported throughout the manuscript were performed using 2.2 mM MgCl_2_ to ensure adequate signal to noise.

In some experiments, plasmid DNA or HIV-1 ERT samples were heat treated prior to incubation in cell-free cGAS assays. For heat treatment experiments, aliquots from ERT reactions were transferred to new PCR tubes and incubated for 20 s in a ProFlex thermocycler at a range of temperatures (37°C–62.5°C). Samples were cooled to room temperature before diluting into cGAS assays. For the time course experiments in Fig 2A, all samples were treated with EFV (1 µM) after ERT to prevent additional reverse transcription that may occur during the cGAS assay at 37°C.

After the cGAS assay, samples were held at 4°C for short term storage or frozen at −80°C. cGAMP levels from cGAS assays were determined by ELISA (Arbor Assays) according to the manufacturers’ instructions. Reaction mixtures were diluted 1:2 in assay buffer and compared to a standard curve using purified 2’3’-cGAMP. Data were analyzed and plotted using GraphPad Prism software. For dose curves comparing responses to LEN, GS-CA1, or PF-74, heat maps were generated using Morpheus (Broad Institute).

### Quantitative PCR analyses

For each ERT reaction, 30 μL of sample was purified using a Qiaquick PCR purification kit (Qiagen) and eluted from the column in 75 μL elution buffer (Qiagen). qPCR reactions were performed and analyzed in triplicate. Primer sets were used to measure early (minus strand strong stop), intermediate (first strand transfer), and late RT (second strand transfer). Standard curves for ERT products were generated by using a 10-fold dilution series of the pSG3Δenv plasmid that were run in parallel with experimental samples and tested using early, intermediate, and late RT primers. Each qPCR sample was comprised of 2.5 μL elution product from purified ERT reactions, 2X SYBR Green Fast qPCR Mix (Abclonal), 1 μL of primers (from a 2 μM working stock), and water to a total of 10 μL reaction volume. qPCR reactions were performed on a QuantStudio 3 (Thermo Fisher) according to the following cycling conditions: 95°C for 10 min, then 45 cycles of 95°C for 10 sec, 55°C for 10 sec, and 72°C for 5 sec, followed by 95°C for 5 sec. A melting curve analysis was then performed going from 65°C to 97°C in 0.11 intervals every 5 sec. Results were analyzed using GraphPad Prism software. For experiments measuring plasmid DNA as a control, plasmid copy numbers determined for each primer set were multiplied by a factor of 1.5 to account for the length difference between the plasmid (14,788 bp) and the reverse transcribed viral genome (∼9,730 bp) in order to plot base-adjusted copy number. At least twice as many copies of early and intermediate products are expected for every one copy of late RT in both the plasmid and HIV-1 full-length genome, based on the number of primer binding sites present in those target sequences.

For quantitation of viral cDNA, 100,000 THP-1 cells were seeded in 96 well plates and infected with HIV-GFP (12.5 nM p24) and polybrene (5 ug/mL) in the absence or presence of EFV (10 µM) and LEN. Infected THP-1s were washed 1X with PBS and DNA was extracted using a DNeasy Blood & Tissue Kit (Qiagen). Reverse transcription products were quantified using primers for early, intermediate (unique to HIV-1-GFP), and late RT, and standard curves were generated using a 10-fold dilution series of the HIV-1-GFPΔenv plasmid run in parallel with experimental samples. Each qPCR sample was comprised of 2.5 μL elution product from viral cDNA extraction, 2X SYBR Green Fast qPCR Mix (Abclonal), 1 μL of primers (from a 2 μM working stock), and water to a total of 10 μL reaction volume. qPCR reactions were performed on a QuantStudio 3 (Thermo Fisher) according to the cycling conditions above and data were analyzed using GraphPad Prism software.

For quantification of host gene expression, 100,000 THP-1 cells were seeded in 96 well plates and infected with HIV-1-GFP (12.5 nM p24) plus polybrene (5 µg/mL) in the absence or presence of 10 nM LEN. Infected THP-1 cells were collected in 1.5 mL tubes at 0 h, 16 h, 24 h, 40 h, and 48 h and washed with 1X PBS. Cells were lysed in RLT Buffer (Qiagen), and RNA was isolated using a RNeasy 96 Kit (Qiagen). Normalized quantities of RNA (50 ng) were converted into cDNA using Superscript III (Thermo Fisher). qPCR reactions were carried out using TaqMan primer probes (Thermo Fisher) and TaqMan Fast Universal PCR Master Mix (Thermo Fisher) on a QuantStudio 3 (Thermo Fisher) in a volume of 10 μL according to the following cycling conditions: 50°C for 2 min, 95°C for 10 min, followed by 40 cycles of 95°C for 15 sec, and 60°C for 1 min. Data were plotted as expression relative to *GAPDH* x 1000.

### Flow cytometry

Infections with HIV-1-GFP reporter viruses were performed in 96 well plates as described previously^19,88^. Briefly, ∼50,000 MDDCs, 25,000 MDMs, or 100,000 THP-1 cells were seeded in 96 well plates and infected as described in the Figure legends with the indicated amounts of virus (typically 12.5 to 50 nM p24) in the presence of polybrene (0.5 – 5 μg/mL). For flow cytometry, 48 h after infection, MDDCs, MDMs, or THP-1 cells were washed with phosphate buffered saline (PBS, Corning) and then exposed to LIVE/DEAD violet (Thermo Fisher) in PBS for 15 min at 4°C in the dark. For intracellular staining using anti-human ISG15-PE (R&D Systems) or SIGLEC1-647 (BD Biosciences), cells were then fixed and permeabilized using a cytofix/cytoperm kit (BD Biosciences) and stained according to the manufacturers’ instructions. Antibody staining was performed for 30 min at room temperature in the dark. Cells were washed and resuspended in PBS with 1% BSA and then analyzed on an Attune flow cytometer (Thermo Fisher). Data were analyzed using FlowJo software (FlowJo LLC). Heat maps were generated using Morpheus (Broad Institute).

### CRISPR-mediated gene editing

THP-1 monocytic cells were modified by lentiCRISPR vectors as previously described^19^. Briefly, we transduced cells in 6-well cluster plates using 800,000 cells per well in 2 mL media with polybrene (5 μg/mL) and concentrated lentiviral stocks (200 μL per well). Two days after transduction, cells were put under selection with puromycin (1 μg/mL, Invivogen) for 4 days and then puromycin was reduced to half-strength for maintenance. Cells were used for experiments beginning 2 weeks after selection. Disruption of target gene expression was confirmed by immunoblot using antibodies to detect cGAS, STING, IRF3, or PQBP1 compared to an actin loading control (as listed in S1 Table).

### Immunoblotting

One million cells were spun down in 1.5 mL tubes and lysed in 150–200 μL SDS sample buffer (2% SDS, 50 mM Tris, 12.5 mM EDTA, plus Halt protease and phosphatase inhibitors (Thermo Fisher)). Samples were sonicated using a microtip Branson sonifier (10–20, 1 s pulses on low output) and protein concentrations were determined using a reducing agent-compatible BCA assay (Thermo Fisher) according to the manufacturer’s instructions. Equal amounts of protein (typically 15–20 μg) were loaded with LDS loading buffer (Life Technologies) and separated on 4–12% gradient poly-acrylamide Bolt Bis-Tris gels (Thermo Fisher). Proteins were transferred to nitrocellulose membranes, blocked using 2.5% BSA (Roche) in Tris buffered saline (TBS) with 0.5% tween. Blots were incubated with primary antibodies, then corresponding fluorescent-conjugated anti-mouse or anti-rabbit secondary antibodies (LI-COR), and then fluorescent images were acquired using an Odyssey Imager (LI-COR). Blot images were analyzed using Image Studio Lite (LI-COR).

## Notes

### Competing Interest Statement

The authors have declared no competing interest.

### Summary of Updates

Manuscript has been revised in response to reviewers' critiques.

## References

1. Zila, V., Margiotta, E., Turonova, B., Muller, T.G., Zimmerli, C.E., Mattei, S., Allegretti, M., Borner, K., Rada, J., Muller, B., et al. (2021). Cone-shaped HIV-1 capsids are transported through intact nuclear pores. Cell 184, 1032–1046 e18. 10.1016/j.cell.2021.01.025.

2. Müller, T.G., Zila, V., Peters, K., Schifferdecker, S., Stanic, M., Lucic, B., Laketa, V., Lusic, M., Müller, B., and Kräusslich, H.-G. (2021). HIV-1 uncoating by release of viral cDNA from capsid-like structures in the nucleus of infected cells. eLife 10, e64776. 10.7554/eLife.64776.

3. Christensen, D.E., Ganser-Pornillos, B.K., Johnson, J.S., Pornillos, O., and Sundquist, W.I. (2020). Reconstitution and visualization of HIV-1 capsid-dependent replication and integration in vitro. Science 370. 10.1126/science.abc8420.

4. Burdick, R.C., Li, C., Munshi, M., Rawson, J.M.O., Nagashima, K., Hu, W.S., and Pathak, V.K. (2020). HIV-1 uncoats in the nucleus near sites of integration. Proc Natl Acad Sci U S A 117, 5486–5493. 10.1073/pnas.1920631117.

5. Li, C., Burdick, R.C., Nagashima, K., Hu, W.-S., and Pathak, V.K. (2021). HIV-1 cores retain their integrity until minutes before uncoating in the nucleus. Proceedings of the National Academy of Sciences 118, e2019467118. 10.1073/pnas.2019467118.

6. Peng, K., Muranyi, W., Glass, B., Laketa, V., Yant, S.R., Tsai, L., Cihlar, T., Muller, B., and Krausslich, H.G. (2014). Quantitative microscopy of functional HIV post-entry complexes reveals association of replication with the viral capsid. Elife 3, e04114. 10.7554/eLife.04114.

7. Bejarano, D.A., Peng, K., Laketa, V., Börner, K., Jost, K.L., Lucic, B., Glass, B., Lusic, M., Müller, B., and Kräusslich, H.-G. (2019). HIV-1 nuclear import in macrophages is regulated by CPSF6-capsid interactions at the nuclear pore complex. Elife 8, e41800. 10.7554/eLife.41800.

8. Schifferdecker, S., Zila, V., Müller, T.G., Sakin, V., Anders-Össwein, M., Laketa, V., Kräusslich, H.-G., and Müller, B. (2022). Direct Capsid Labeling of Infectious HIV-1 by Genetic Code Expansion Allows Detection of Largely Complete Nuclear Capsids and Suggests Nuclear Entry of HIV-1 Complexes via Common Routes. mBio 13, e0195922. 10.1128/mbio.01959-22.

9. Fu, L., Weiskopf, E.N., Akkermans, O., Swanson, N.A., Cheng, S., Schwartz, T.U., and Görlich, D. (2024). HIV-1 capsids enter the FG phase of nuclear pores like a transport receptor. Nature 626, 843–851. 10.1038/s41586-023-06966-w.

10. Dickson, C.F., Hertel, S., Tuckwell, A.J., Li, N., Ruan, J., Al-Izzi, S.C., Ariotti, N., Sierecki, E., Gambin, Y., Morris, R.G., et al. (2024). The HIV capsid mimics karyopherin engagement of FG-nucleoporins. Nature 626, 836–842. 10.1038/s41586-023-06969-7.

11. Francis, A.C., Marin, M., Singh, P.K., Achuthan, V., Prellberg, M.J., Palermino-Rowland, K., Lan, S., Tedbury, P.R., Sarafianos, S.G., Engelman, A.N., et al. (2020). HIV-1 replication complexes accumulate in nuclear speckles and integrate into speckle-associated genomic domains. Nat Commun 11, 3505. 10.1038/s41467-020-17256-8.

12. Dharan, A., Bachmann, N., Talley, S., Zwikelmaier, V., and Campbell, E.M. (2020). Nuclear pore blockade reveals that HIV-1 completes reverse transcription and uncoating in the nucleus. Nat Microbiol 5, 1088–1095. 10.1038/s41564-020-0735-8.

13. Rensen, E., Mueller, F., Scoca, V., Parmar, J.J., Souque, P., Zimmer, C., and Di Nunzio, F. (2021). Clustering and reverse transcription of HIV-1 genomes in nuclear niches of macrophages. The EMBO Journal 40, e105247. 10.15252/embj.2020105247.

14. Selyutina, A., Persaud, M., Lee, K., KewalRamani, V., and Diaz-Griffero, F. (2020). Nuclear Import of the HIV-1 Core Precedes Reverse Transcription and Uncoating. Cell Rep 32, 108201. 10.1016/j.celrep.2020.108201.

15. Sumner, R.P., Harrison, L., Touizer, E., Peacock, T.P., Spencer, M., Zuliani-Alvarez, L., and Towers, G.J. (2020). Disrupting HIV-1 capsid formation causes cGAS sensing of viral DNA. EMBO J 39, e103958. 10.15252/embj.2019103958.

16. Rasaiyaah, J., Tan, C.P., Fletcher, A.J., Price, A.J., Blondeau, C., Hilditch, L., Jacques, D.A., Selwood, D.L., James, L.C., Noursadeghi, M., et al. (2013). HIV-1 evades innate immune recognition through specific cofactor recruitment. Nature 503, 402–405. 10.1038/nature12769.

17. Lahaye, X., Satoh, T., Gentili, M., Cerboni, S., Conrad, C., Hurbain, I., El Marjou, A., Lacabaratz, C., Lelievre, J.D., and Manel, N. (2013). The capsids of HIV-1 and HIV-2 determine immune detection of the viral cDNA by the innate sensor cGAS in dendritic cells. Immunity 39, 1132–1142. 10.1016/j.immuni.2013.11.002.

18. Gao, D., Wu, J., Wu, Y.T., Du, F., Aroh, C., Yan, N., Sun, L., and Chen, Z.J. (2013). Cyclic GMP-AMP synthase is an innate immune sensor of HIV and other retroviruses. Science 341, 903–906. 10.1126/science.1240933.

19. Johnson, J.S., Lucas, S.Y., Amon, L.M., Skelton, S., Nazitto, R., Carbonetti, S., Sather, D.N., Littman, D.R., and Aderem, A. (2018). Reshaping of the Dendritic Cell Chromatin Landscape and Interferon Pathways during HIV Infection. Cell Host Microbe 23, 366–381 e9. 10.1016/j.chom.2018.01.012.

20. Yan, N., Regalado-Magdos, A.D., Stiggelbout, B., Lee-Kirsch, M.A., and Lieberman, J. (2010). The cytosolic exonuclease TREX1 inhibits the innate immune response to human immunodeficiency virus type 1. Nat Immunol 11, 1005–1013. 10.1038/ni.1941.

21. Manel, N., Hogstad, B., Wang, Y., Levy, D.E., Unutmaz, D., and Littman, D.R. (2010). A cryptic sensor for HIV-1 activates antiviral innate immunity in dendritic cells. Nature 467, 214–217. 10.1038/nature09337.

22. Yum, S., Li, M., Fang, Y., and Chen, Z.J. (2021). TBK1 recruitment to STING activates both IRF3 and NF-κB that mediate immune defense against tumors and viral infections. Proceedings of the National Academy of Sciences 118, e2100225118. 10.1073/pnas.2100225118.

23. Liu, S., Cai, X., Wu, J., Cong, Q., Chen, X., Li, T., Du, F., Ren, J., Wu, Y.T., Grishin, N.V., et al. (2015). Phosphorylation of innate immune adaptor proteins MAVS, STING, and TRIF induces IRF3 activation. Science 347, aaa2630. 10.1126/science.aaa2630.

24. Schoggins, J.W., Wilson, S.J., Panis, M., Murphy, M.Y., Jones, C.T., Bieniasz, P., and Rice, C.M. (2011). A diverse range of gene products are effectors of the type I interferon antiviral response. Nature 472, 481–485. 10.1038/nature09907.

25. Johnson, J.S., De Veaux, N., Rives, A.W., Lahaye, X., Lucas, S.Y., Perot, B.P., Luka, M., Garcia-Paredes, V., Amon, L.M., Watters, A., et al. (2020). A Comprehensive Map of the Monocyte-Derived Dendritic Cell Transcriptional Network Engaged upon Innate Sensing of HIV. Cell Rep 30, 914–931 e9. 10.1016/j.celrep.2019.12.054.

26. Kane, M., Zang, T.M., Rihn, S.J., Zhang, F., Kueck, T., Alim, M., Schoggins, J., Rice, C.M., Wilson, S.J., and Bieniasz, P.D. (2016). Identification of Interferon-Stimulated Genes with Antiretroviral Activity. Cell Host Microbe 20, 392–405. 10.1016/j.chom.2016.08.005.

27. OhAinle, M., Helms, L., Vermeire, J., Roesch, F., Humes, D., Basom, R., Delrow, J.J., Overbaugh, J., and Emerman, M. (2018). A virus-packageable CRISPR screen identifies host factors mediating interferon inhibition of HIV. Elife 7. 10.7554/eLife.39823.

28. Siddiqui, M.A., Saito, A., Halambage, U.D., Ferhadian, D., Fischer, D.K., Francis, A.C., Melikyan, G.B., Ambrose, Z., Aiken, C., and Yamashita, M. (2019). A Novel Phenotype Links HIV-1 Capsid Stability to cGAS-Mediated DNA Sensing. J Virol 93. 10.1128/JVI.00706-19.

29. Papa, G., Albecka, A., Mallery, D., Vaysburd, M., Renner, N., and James, L.C. (2023). IP6-stabilised HIV capsids evade cGAS/STING-mediated host immune sensing. EMBO reports 24, e56275. 10.15252/embr.202256275.

30. Zuliani-Alvarez, L., Govasli, M.L., Rasaiyaah, J., Monit, C., Perry, S.O., Sumner, R.P., McAlpine-Scott, S., Dickson, C., Rifat Faysal, K.M., Hilditch, L., et al. (2022). Evasion of cGAS and TRIM5 defines pandemic HIV. Nat Microbiol 7, 1762–1776. 10.1038/s41564-022-01247-0.

31. Lahaye, X., Gentili, M., Silvin, A., Conrad, C., Picard, L., Jouve, M., Zueva, E., Maurin, M., Nadalin, F., Knott, G.J., et al. (2018). NONO Detects the Nuclear HIV Capsid to Promote cGAS-Mediated Innate Immune Activation. Cell 175, 488–501 e22. 10.1016/j.cell.2018.08.062.

32. Yoh, S.M., Mamede, J.I., Lau, D., Ahn, N., Sánchez-Aparicio, M.T., Temple, J., Tuckwell, A., Fuchs, N.V., Cianci, G.C., Riva, L., et al. (2022). Recognition of HIV-1 capsid by PQBP1 licenses an innate immune sensing of nascent HIV-1 DNA. Mol Cell 82, 2871–2884.e6. 10.1016/j.molcel.2022.06.010.

33. Eschbach, J.E., Puray-Chavez, M., Mohammed, S., Wang, Q., Xia, M., Huang, L.-C., Shan, L., and Kutluay, S.B. (2024). HIV-1 capsid stability and reverse transcription are finely balanced to minimize sensing of reverse transcription products via the cGAS-STING pathway. mBio 0, e00348–24. 10.1128/mbio.00348-24.

34. Dick, R.A., Zadrozny, K.K., Xu, C., Schur, F.K.M., Lyddon, T.D., Ricana, C.L., Wagner, J.M., Perilla, J.R., Ganser-Pornillos, B.K., Johnson, M.C., et al. (2018). Inositol phosphates are assembly co-factors for HIV-1. Nature 560, 509–512. 10.1038/s41586-018-0396-4.

35. Mallery, D.L., Márquez, C.L., McEwan, W.A., Dickson, C.F., Jacques, D.A., Anandapadamanaban, M., Bichel, K., Towers, G.J., Saiardi, A., Böcking, T., et al. (2018). IP6 is an HIV pocket factor that prevents capsid collapse and promotes DNA synthesis. Elife 7, e35335. 10.7554/eLife.35335.

36. Jennings, J., Shi, J., Varadarajan, J., Jamieson, P.J., and Aiken, C. (2020). The Host Cell Metabolite Inositol Hexakisphosphate Promotes Efficient Endogenous HIV-1 Reverse Transcription by Stabilizing the Viral Capsid. mBio 11, 10.1128/mbio.02820-20. 10.1128/mbio.02820-20.

37. Mackenzie, K.J., Carroll, P., Martin, C.-A., Murina, O., Fluteau, A., Simpson, D.J., Olova, N., Sutcliffe, H., Rainger, J.K., Leitch, A., et al. (2017). cGAS surveillance of micronuclei links genome instability to innate immunity. Nature 548, 461–465. 10.1038/nature23449.

38. Forshey, B.M., von Schwedler, U., Sundquist, W.I., and Aiken, C. (2002). Formation of a human immunodeficiency virus type 1 core of optimal stability is crucial for viral replication. J Virol 76, 5667–5677. 10.1128/jvi.76.11.5667-5677.2002.

39. von Schwedler, U.K., Stray, K.M., Garrus, J.E., and Sundquist, W.I. (2003). Functional Surfaces of the Human Immunodeficiency Virus Type 1 Capsid Protein. Journal of Virology 77, 5439–5450. 10.1128/jvi.77.9.5439-5450.2003.

40. Ganser-Pornillos, B.K., von Schwedler, U.K., Stray, K.M., Aiken, C., and Sundquist, W.I. (2004). Assembly Properties of the Human Immunodeficiency Virus Type 1 CA Protein. Journal of Virology 78, 2545–2552. 10.1128/jvi.78.5.2545-2552.2004.

41. Sowd, G.A., Shi, J., Fulmer, A., and Aiken, C. (2023). HIV-1 capsid stability enables inositol phosphate-independent infection of target cells and promotes integration into genes. PLOS Pathogens 19, e1011423. 10.1371/journal.ppat.1011423.

42. Kleinpeter, A.B., Zhu, Y., Mallery, D.L., Ablan, S.D., Chen, L., Hardenbrook, N., Saiardi, A., James, L.C., Zhang, P., and Freed, E.O. (2023). The Effect of Inositol Hexakisphosphate on HIV-1 Particle Production and Infectivity can be Modulated by Mutations that Affect the Stability of the Immature Gag Lattice. Journal of Molecular Biology 435, 168037. 10.1016/j.jmb.2023.168037.

43. Letcher, A.J., Schell, M.J., and Irvine, R.F. (2008). Do mammals make all their own inositol hexakisphosphate? Biochem J 416, 263–270. 10.1042/BJ20081417.

44. Kumar, S., Morrison, J.H., Dingli, D., and Poeschla, E. (2018). HIV-1 Activation of Innate Immunity Depends Strongly on the Intracellular Level of TREX1 and Sensing of Incomplete Reverse Transcription Products. J Virol 92. 10.1128/JVI.00001-18.

45. Yant, S.R., Mulato, A., Hansen, D., Tse, W.C., Niedziela-Majka, A., Zhang, J.R., Stepan, G.J., Jin, D., Wong, M.H., Perreira, J.M., et al. (2019). A highly potent long-acting small-molecule HIV-1 capsid inhibitor with efficacy in a humanized mouse model. Nat Med 25, 1377–1384. 10.1038/s41591-019-0560-x.

46. Link, J.O., Rhee, M.S., Tse, W.C., Zheng, J., Somoza, J.R., Rowe, W., Begley, R., Chiu, A., Mulato, A., Hansen, D., et al. (2020). Clinical targeting of HIV capsid protein with a long-acting small molecule. Nature 584, 614–618. 10.1038/s41586-020-2443-1.

47. Deshpande, A., Bryer, A.J., Andino-Moncada, J.R., Shi, J., Hong, J., Torres, C., Harel, S., Francis, A.C., Perilla, J.R., Aiken, C., et al. (2024). Elasticity of the HIV-1 core facilitates nuclear entry and infection. PLoS Pathog 20, e1012537. 10.1371/journal.ppat.1012537.

48. Highland, C.M., Tan, A., Ricaña, C.L., Briggs, J.A.G., and Dick, R.A. (2023). Structural insights into HIV-1 polyanion-dependent capsid lattice formation revealed by single particle cryo-EM. Proc Natl Acad Sci U S A 120, e2220545120. 10.1073/pnas.2220545120.

49. Bester, S.M., Wei, G., Zhao, H., Adu-Ampratwum, D., Iqbal, N., Courouble, V.V., Francis, A.C., Annamalai, A.S., Singh, P.K., Shkriabai, N., et al. (2020). Structural and mechanistic bases for a potent HIV-1 capsid inhibitor. Science 370, 360–364. 10.1126/science.abb4808.

50. Faysal, K.R., Walsh, J.C., Renner, N., Márquez, C.L., Shah, V.B., Tuckwell, A.J., Christie, M.P., Parker, M.W., Turville, S.G., Towers, G.J., et al. (2024). Pharmacologic hyperstabilisation of the HIV-1 capsid lattice induces capsid failure. eLife 13, e83605. 10.7554/eLife.83605.

51. Wang, C., Guan, Y., Lv, M., Zhang, R., Guo, Z., Wei, X., Du, X., Yang, J., Li, T., Wan, Y., et al. (2018). Manganese Increases the Sensitivity of the cGAS-STING Pathway for Double-Stranded DNA and Is Required for the Host Defense against DNA Viruses. Immunity 48, 675–687 e7. 10.1016/j.immuni.2018.03.017.

52. Blair, W.S., Pickford, C., Irving, S.L., Brown, D.G., Anderson, M., Bazin, R., Cao, J., Ciaramella, G., Isaacson, J., Jackson, L., et al. (2010). HIV Capsid is a Tractable Target for Small Molecule Therapeutic Intervention. PLoS Pathog 6, e1001220. 10.1371/journal.ppat.1001220.

53. Bhattacharya, A., Alam, S.L., Fricke, T., Zadrozny, K., Sedzicki, J., Taylor, A.B., Demeler, B., Pornillos, O., Ganser-Pornillos, B.K., Diaz-Griffero, F., et al. (2014). Structural basis of HIV-1 capsid recognition by PF74 and CPSF6. Proceedings of the National Academy of Sciences 111, 18625–18630. 10.1073/pnas.1419945112.

54. Segal-Maurer, S., DeJesus, E., Stellbrink, H.-J., Castagna, A., Richmond, G.J., Sinclair, G.I., Siripassorn, K., Ruane, P.J., Berhe, M., Wang, H., et al. (2022). Capsid Inhibition with Lenacapavir in Multidrug-Resistant HIV-1 Infection. N Engl J Med 386, 1793–1803. 10.1056/NEJMoa2115542.

55. Schirra, R.T., dos Santos, N.F.B., Zadrozny, K.K., Kucharska, I., Ganser-Pornillos, B.K., and Pornillos, O. (2023). A molecular switch modulates assembly and host factor binding of the HIV-1 capsid. Nat Struct Mol Biol 30, 383–390. 10.1038/s41594-022-00913-5.

56. Sanjana, N.E., Shalem, O., and Zhang, F. (2014). Improved vectors and genome-wide libraries for CRISPR screening. Nat Methods 11, 783–784. 10.1038/nmeth.3047.

57. Yoh, S.M., Schneider, M., Seifried, J., Soonthornvacharin, S., Akleh, R.E., Olivieri, K.C., De Jesus, P.D., Ruan, C., de Castro, E., Ruiz, P.A., et al. (2015). PQBP1 Is a Proximal Sensor of the cGAS-Dependent Innate Response to HIV-1. Cell 161, 1293–1305. 10.1016/j.cell.2015.04.050.

58. Hrecka, K., Hao, C., Gierszewska, M., Swanson, S.K., Kesik-Brodacka, M., Srivastava, S., Florens, L., Washburn, M.P., and Skowronski, J. (2011). Vpx relieves inhibition of HIV-1 infection of macrophages mediated by the SAMHD1 protein. Nature 474, 658–661. 10.1038/nature10195.

59. Laguette, N., Sobhian, B., Casartelli, N., Ringeard, M., Chable-Bessia, C., Segeral, E., Yatim, A., Emiliani, S., Schwartz, O., and Benkirane, M. (2011). SAMHD1 is the dendritic- and myeloid-cell-specific HIV-1 restriction factor counteracted by Vpx. Nature 474, 654–657. 10.1038/nature10117.

60. McCauley, S.M., Kim, K., Nowosielska, A., Dauphin, A., Yurkovetskiy, L., Diehl, W.E., and Luban, J. (2018). Intron-containing RNA from the HIV-1 provirus activates type I interferon and inflammatory cytokines. Nat Commun 9, 5305. 10.1038/s41467-018-07753-2.

61. Akiyama, H., Miller, C.M., Ettinger, C.R., Belkina, A.C., Snyder-Cappione, J.E., and Gummuluru, S. (2018). HIV-1 intron-containing RNA expression induces innate immune activation and T cell dysfunction. Nat Commun 9, 3450. 10.1038/s41467-018-05899-7.

62. Gondim, M.V.P., Sherrill-Mix, S., Bibollet-Ruche, F., Russell, R.M., Trimboli, S., Smith, A.G., Li, Y., Liu, W., Avitto, A.N., DeVoto, J.C., et al. (2021). Heightened resistance to host type 1 interferons characterizes HIV-1 at transmission and after antiretroviral therapy interruption. Sci Transl Med 13. 10.1126/scitranslmed.abd8179.

63. Iyer, S.S., Bibollet-Ruche, F., Sherrill-Mix, S., Learn, G.H., Plenderleith, L., Smith, A.G., Barbian, H.J., Russell, R.M., Gondim, M.V., Bahari, C.Y., et al. (2017). Resistance to type 1 interferons is a major determinant of HIV-1 transmission fitness. Proc Natl Acad Sci U S A 114, E590–E599. 10.1073/pnas.1620144114.

64. Beignon, A.S., McKenna, K., Skoberne, M., Manches, O., DaSilva, I., Kavanagh, D.G., Larsson, M., Gorelick, R.J., Lifson, J.D., and Bhardwaj, N. (2005). Endocytosis of HIV-1 activates plasmacytoid dendritic cells via Toll-like receptor-viral RNA interactions. The Journal of clinical investigation 115, 3265–3275. 10.1172/JCI26032.

65. Lepelley, A., Louis, S., Sourisseau, M., Law, H.K., Pothlichet, J., Schilte, C., Chaperot, L., Plumas, J., Randall, R.E., Si-Tahar, M., et al. (2011). Innate sensing of HIV-infected cells. PLoS Pathog 7, e1001284. 10.1371/journal.ppat.1001284.

66. Wang, Q., Gao, H., Clark, K.M., Mugisha, C.S., Davis, K., Tang, J.P., Harlan, G.H., DeSelm, C.J., Presti, R.M., Kutluay, S.B., et al. (2021). CARD8 is an inflammasome sensor for HIV-1 protease activity. Science 371. 10.1126/science.abe1707.

67. Monroe, K.M., Yang, Z., Johnson, J.R., Geng, X., Doitsh, G., Krogan, N.J., and Greene, W.C. (2014). IFI16 DNA sensor is required for death of lymphoid CD4 T cells abortively infected with HIV. Science 343, 428–432. 10.1126/science.1243640.

68. Doitsh, G., Cavrois, M., Lassen, K.G., Zepeda, O., Yang, Z., Santiago, M.L., Hebbeler, A.M., and Greene, W.C. (2010). Abortive HIV Infection Mediates CD4 T-Cell Depletion and Inflammation in Human Lymphoid Tissue. Cell 143, 789–801. 10.1016/j.cell.2010.11.001.

69. Martin-Gayo, E., Buzon, M.J., Ouyang, Z., Hickman, T., Cronin, J., Pimenova, D., Walker, B.D., Lichterfeld, M., and Yu, X.G. (2015). Potent Cell-Intrinsic Immune Responses in Dendritic Cells Facilitate HIV-1-Specific T Cell Immunity in HIV-1 Elite Controllers. PLoS Pathog 11, e1004930. 10.1371/journal.ppat.1004930.

70. Martin-Gayo, E., Cole, M.B., Kolb, K.E., Ouyang, Z., Cronin, J., Kazer, S.W., Ordovas-Montanes, J., Lichterfeld, M., Walker, B.D., Yosef, N., et al. (2018). A Reproducibility-Based Computational Framework Identifies an Inducible, Enhanced Antiviral State in Dendritic Cells from HIV-1 Elite Controllers. Genome Biol 19, 10. 10.1186/s13059-017-1385-x.

71. Migueles, S.A., Laborico, A.C., Shupert, W.L., Sabbaghian, M.S., Rabin, R., Hallahan, C.W., Van Baarle, D., Kostense, S., Miedema, F., McLaughlin, M., et al. (2002). HIV-specific CD8+ T cell proliferation is coupled to perforin expression and is maintained in nonprogressors. Nat Immunol 3, 1061–1068. 10.1038/ni845.

72. Ndhlovu, Z.M., Kamya, P., Mewalal, N., Kloverpris, H.N., Nkosi, T., Pretorius, K., Laher, F., Ogunshola, F., Chopera, D., Shekhar, K., et al. (2015). Magnitude and Kinetics of CD8+ T Cell Activation during Hyperacute HIV Infection Impact Viral Set Point. Immunity 43, 591– 604. 10.1016/j.immuni.2015.08.012.

73. Jang, S., and Engelman, A.N. (2023). Capsid-host interactions for HIV-1 ingress. Microbiol Mol Biol Rev 87, e0004822. 10.1128/mmbr.00048-22.

74. Zhuang, S., and Torbett, B.E. (2021). Interactions of HIV-1 Capsid with Host Factors and Their Implications for Developing Novel Therapeutics. Viruses 13, 417. 10.3390/v13030417.

75. Luban, J., Bossolt, K.L., Franke, E.K., Kalpana, G.V., and Goff, S.P. (1993). Human immunodeficiency virus type 1 Gag protein binds to cyclophilins A and B. Cell 73, 1067– 1078. 10.1016/0092-8674(93)90637-6.

76. Lee, K., Ambrose, Z., Martin, T.D., Oztop, I., Mulky, A., Julias, J.G., Vandegraaff, N., Baumann, J.G., Wang, R., Yuen, W., et al. (2010). Flexible use of nuclear import pathways by HIV-1. Cell Host Microbe 7, 221–233. 10.1016/j.chom.2010.02.007.

77. Zhong, Z., Ning, J., Boggs, E.A., Jang, S., Wallace, C., Telmer, C., Bruchez, M.P., Ahn, J., Engelman, A.N., Zhang, P., et al. (2021). Cytoplasmic CPSF6 Regulates HIV-1 Capsid Trafficking and Infection in a Cyclophilin A-Dependent Manner. mBio 12, e03142–20. 10.1128/mBio.03142-20.

78. De Iaco, A., and Luban, J. (2014). Cyclophilin A promotes HIV-1 reverse transcription but its effect on transduction correlates best with its effect on nuclear entry of viral cDNA. Retrovirology 11, 11. 10.1186/1742-4690-11-11.

79. Chin, C.R., Perreira, J.M., Savidis, G., Portmann, J.M., Aker, A.M., Feeley, E.M., Smith, M.C., and Brass, A.L. (2015). Direct Visualization of HIV-1 Replication Intermediates Shows that Capsid and CPSF6 Modulate HIV-1 Intra-nuclear Invasion and Integration. Cell Rep 13, 1717–1731. 10.1016/j.celrep.2015.10.036.

80. Schaller, T., Ocwieja, K.E., Rasaiyaah, J., Price, A.J., Brady, T.L., Roth, S.L., Hué, S., Fletcher, A.J., Lee, K., KewalRamani, V.N., et al. (2011). HIV-1 capsid-cyclophilin interactions determine nuclear import pathway, integration targeting and replication efficiency. PLoS Pathog 7, e1002439. 10.1371/journal.ppat.1002439.

81. Achuthan, V., Perreira, J.M., Sowd, G.A., Puray-Chavez, M., McDougall, W.M., Paulucci-Holthauzen, A., Wu, X., Fadel, H.J., Poeschla, E.M., Multani, A.S., et al. (2018). Capsid-CPSF6 Interaction Licenses Nuclear HIV-1 Trafficking to Sites of Viral DNA Integration. Cell Host Microbe 24, 392–404 e8. 10.1016/j.chom.2018.08.002.

82. Li, Y.-L., Chandrasekaran, V., Carter, S.D., Woodward, C.L., Christensen, D.E., Dryden, K.A., Pornillos, O., Yeager, M., Ganser-Pornillos, B.K., Jensen, G.J., et al. (2016). Primate TRIM5 proteins form hexagonal nets on HIV-1 capsids. Elife 5, e16269. 10.7554/eLife.16269.

83. Stremlau, M., Owens, C.M., Perron, M.J., Kiessling, M., Autissier, P., and Sodroski, J. (2004). The cytoplasmic body component TRIM5alpha restricts HIV-1 infection in Old World monkeys. Nature 427, 848–853. 10.1038/nature02343.

84. Stremlau, M., Perron, M., Lee, M., Li, Y., Song, B., Javanbakht, H., Diaz-Griffero, F., Anderson, D.J., Sundquist, W.I., and Sodroski, J. (2006). Specific recognition and accelerated uncoating of retroviral capsids by the TRIM5α restriction factor. Proceedings of the National Academy of Sciences 103, 5514–5519. 10.1073/pnas.0509996103.

85. Pertel, T., Hausmann, S., Morger, D., Zuger, S., Guerra, J., Lascano, J., Reinhard, C., Santoni, F.A., Uchil, P.D., Chatel, L., et al. (2011). TRIM5 is an innate immune sensor for the retrovirus capsid lattice. Nature 472, 361–365. 10.1038/nature09976.

86. Jeremiah, N., Ferran, H., Antoniadou, K., De Azevedo, K., Nikolic, J., Maurin, M., Benaroch, P., and Manel, N. (2023). RELA tunes innate-like interferon I/III responses in human T cells. J Exp Med 220, e20220666. 10.1084/jem.20220666.

87. Elsner, C., Ponnurangam, A., Kazmierski, J., Zillinger, T., Jansen, J., Todt, D., Dohner, K., Xu, S., Ducroux, A., Kriedemann, N., et al. (2020). Absence of cGAS-mediated type I IFN responses in HIV-1-infected T cells. Proc Natl Acad Sci U S A 117, 19475–19486. 10.1073/pnas.2002481117.

88. Greiner, D., Scott, T.M., Olson, G.S., Aderem, A., Roh-Johnson, M., and Johnson, J.S. (2022). Genetic Modification of Primary Human Myeloid Cells to Study Cell Migration, Activation, and Organelle Dynamics. Curr Protoc 2, e514. 10.1002/cpz1.514.

89. Mangeot, P.E., Nègre, D., Dubois, B., Winter, A.J., Leissner, P., Mehtali, M., Kaiserlian, D., Cosset, F.L., and Darlix, J.L. (2000). Development of minimal lentivirus vectors derived from simian immunodeficiency virus (SIVmac251) and their use for gene transfer into human dendritic cells. J. Virol. 74, 8307–8315.

90. Ott, D.E. (2009). Purification of HIV-1 Virions by Subtilisin Digestion or CD45 Immunoaffinity Depletion for Biochemical Studies. In HIV Protocols Methods In Molecular Biology^TM^., V. R. Prasad and G. V. Kalpana, eds. (Humana Press), pp. 15–25. 10.1007/978-1-59745-170-3_2.

91. Zhou, W., Whiteley, A.T., de Oliveira Mann, C.C., Morehouse, B.R., Nowak, R.P., Fischer, E.S., Gray, N.S., Mekalanos, J.J., and Kranzusch, P.J. (2018). Structure of the Human cGAS–DNA Complex Reveals Enhanced Control of Immune Surveillance. Cell 174, 300–311.e11. 10.1016/j.cell.2018.06.026.

92. Zhou, W., Whiteley, A.T., and Kranzusch, P.J. (2019). Chapter Two - Analysis of human cGAS activity and structure. In Methods in Enzymology DNA Sensors and Inflammasomes., J. Sohn, ed. (Academic Press), pp. 13–40. 10.1016/bs.mie.2019.04.012.

93. Kennedy, E.M., Gavegnano, C., Nguyen, L., Slater, R., Lucas, A., Fromentin, E., Schinazi, R.F., and Kim, B. (2010). Ribonucleoside triphosphates as substrate of human immunodeficiency virus type 1 reverse transcriptase in human macrophages. J Biol Chem 285, 39380–39391. 10.1074/jbc.M110.178582.

94. Diamond, T.L., Roshal, M., Jamburuthugoda, V.K., Reynolds, H.M., Merriam, A.R., Lee, K.Y., Balakrishnan, M., Bambara, R.A., Planelles, V., Dewhurst, S., et al. (2004). Macrophage tropism of HIV-1 depends on efficient cellular dNTP utilization by reverse transcriptase. J Biol Chem 279, 51545–51553. 10.1074/jbc.M408573200.

