## Supplementary information for "Cell-free assays reveal that the HIV-1 capsid protects reverse transcripts from cGAS"

[†]Contributed equally

**Supplementary Information includes:**

**S1 Table. Reagents used in this study.**

**S1 Fig. Capsid stability can be affected by heat or CA mutations.**

**S2 Fig. Cell-free sensing of HIV-1 requires virion permeabilization, magnesium, NTPs, and cGAS.**

**S3 Fig. Specific CA mutations confer resistance to LEN.**

**S4 Fig. LEN augments innate immune responses across a range of virus doses.**

**S5 Fig. PQBP1 does not affect cGAS activity *in vitro* and is not essential in cells.**

**S1 Movie. Model for LEN-mediated disruption of HIV-1 cores and cGAS-STING activation. (separate file)**

A molecular animation depicting an HIV-1 viral core in the cytoplasm of myeloid cells, with reverse transcription proceeding inside an intact or largely intact capsid. In the presence of lenacapavir, accumulated reverse transcripts can become exposed by drug-induced capsid rupturing, enabling viral DNA detection and downstream activation of interferon production via the cGAS-STING-IRF3 pathway.

**S1 Table. Reagents used in this study.**

| REAGENT | Source | Catalogue # |
|---|---|---|
| Antibodies | | |
| cGAS | Cell Signaling | Cat# 79978<br>RRID:AB_2905508 |
| STING | Cell Signaling | Cat# 13647<br>RRID:AB_2732796 |
| IRF3 | Cell Signaling | Cat# 11904<br>RRID:AB_2722521 |
| PQBP1 | Proteintech | Cat# 16264-1-AP<br>RRID:AB_10792928 |
| ISG15-PE | R&D Systems | Cat# IC8044P |
| SIGLEC1-647 | BD Biosciences | Cat# 565295<br>RRID:AB_2739163 |
| Actin | BD Biosciences | Cat# 612656<br>RRID:AB_2289199 |
| IRDye 680RD Donkey anti-Mouse fluorescent antibody | LI-COR | Cat# 926-68072<br>RRID:AB_10953628 |
| IRDye 800CW Goat anti-Rabbit fluorescent antibody | LI-COR | Cat# 926-32211<br>RRID:AB_621843 |
| Bacterial and virus strains | | |
| Stbl3 competent *E. coli* | Thermo Fisher | Cat# C737303 |
| BL21-CodonPlus (DE3)-RIPL *E. coli* | Agilent | Cat# 230280 |
| BL21-CodonPlus (DE3)-RIL *E. coli* | Agilent | Cat# 230245 |
| HIV-1 SG3ΔENV | NIH AIDS Reagent Program | Addgene plasmid #11051 |
| HIV-1 SG3ΔENV RT D185A | Christensen et al., 2020 | Addgene plasmid #145782 |
| HIV-1 SG3ΔENV CA E45A | Christensen et al., 2020 | Addgene plasmid #145783 |
| HIV-1 SG3ΔENV CA Q63/67A | Christensen et al., 2020 | Addgene plasmid #145784 |
| HIV-1 SG3ΔENV CA M66I | Christensen et al., 2020 | Addgene plasmid #149687 |
| HIV-1 SG3ΔENV CA Q67H/N74D | This study | Addgene plasmid #217442 |
| HIV-1-GFPΔENV | This study | Addgene plasmid #217437 |
| HIV-1-GFP CA E45A | This study | Addgene plasmid #217438 |
| HIV-1-GFP CA Q63/67A | This study | Addgene plasmid #217439 |
| HIV-1-GFP CA M66I | This study | Addgene plasmid #217440 |
| HIV-1-GFP CA Q67H/N74D | This study | Addgene plasmid #217441 |
| pSIV3+ | Mangeot et al., 2000 | N/A |

| | | |
|---|---|---|
| Chemicals, peptides, and recombinant proteins | | |
| Melittin | Sigma Aldrich | Cat# M2272-5MG |
| IP6 | Sigma Aldrich | Cat# I5125-50G |
| rATP | Promega | Cat# E6011 |
| rGTP | Promega | Cat# E6031 |
| rCTP | Promega | Cat# E6041 |
| rUTP | Promega | Cat# E6021 |
| dATP | Promega | Cat# U120D |
| dGTP | Promega | Cat# U121D |
| dCTP | Promega | Cat# U122D |
| dTTP | Promega | Cat# U123D |
| HEPES | Gibco | Cat# 15630080 |
| Phosphate Buffer Saline (PBS) | Gibco | Cat# 14190250 |
| EDTA 0.5M | Invitrogen | Cat# AM9260G |
| Penicillin-Streptomycin | Thermo Fisher | Cat# 15140-122 |
| Chloramphenicol | Thermo Fisher | Cat# B20841.22 |
| 2-Mercaptoethanol | Thermo Fisher | Cat# 21985023 |
| MEM non-essential amino acids | Thermo Fisher | Cat# 11140050 |
| RPMI 1640 medium | Thermo Fisher | Cat# 11875-119 |
| DMEM | Thermo Fisher | Cat# 11995073 |
| Ficoll-Paque Plus | GE Healthcare | Cat# 17-1440-02 |
| Fetal Bovine Serum | Gibco | (Lot #1982147) |
| Bovine Serum Albumin Fraction V, heat shock | Sigma Aldrich | Cat# 03116956001 |
| Recombinant Human IL-4 | Miltenyi Biotech | Cat# 130-093-922 |
| Recombinant Human GM-CSF | Miltenyi Biotech | Cat# 130-093-867 |
| poly-L-lysine hydrobromide | MP Biomedicals | Cat# 0219454405 |
| Polyethylenimine “Max”, (Mw 40,000) - High Potency Linear PEI | Polysciences, Inc. | Cat# 24765-1 |
| Paraformaldehyde | Electron Microscopy Sciences | Cat# 15713-S |
| Polybrene | Sigma | Cat# TR-1003-G |
| Puromycin | Invivogen | Cat# ant-pr-1 |
| Halt protease and phosphatase inhibitor | Thermo Fisher | Cat# 78441 |
| Efavirenz | Selleck Chemicals | S4685 |
| Elvitegravir | Selleck Chemicals | S2001 |
| Lenacapavir | MCE | HY-111964 |
| GS-CA1 | Gilead | N/A |
| PF-74 (PF-3450074) | MCE | HY-120072 |
| BOLT transfer buffer (20X) | Invitrogen | BT00061 |
| Novex NuPAGE MOPS SDS Running Buffer (20X) | Invitrogen | NP0001 |
| NuPAGE LDS Sample Buffer (4X) | Invitrogen | NP0007 |
| Benzonase | Sigma | Cat# E1014 |
| Subtilisin Protease from Bacillus | Sigma | Cat# P5380 |
| PMSF | Roche | Cat# 10837091001 |
| NaCl | Thermo Fisher | Cat# BP358-10 |
| Tris-HCl | Thermo Fisher | Cat# 228030051 |
| Tris-base | Thermo Fisher | Cat# BP152-500 |
| $MgCl_2$ | Thermo Fisher | Cat# 41341-5000 |
| Isopropyl ß-D-1-thiogalactopyranoside | GoldBio | Cat# I2481C50 |
| 2xYT media | Sigma | Cat# Y1003 |

| | | |
|---|---|---|
| Imidazole | Thermo Fisher | Cat# AC122020020 |
| Lysozyme | GoldBio | Cat# L-040-100 |
| Pepstatin | Sigma | Cat #11524488001 |
| Leupeptin | Sigma | Cat #L2884 |
| Aprotinin | Sigma | Cat# 10981532001 |
| DNase I | GoldBio | Cat# D-300-500 |
| $His_6$-ULP1 protease | Produced in-house | Lao et al. 2018, J. Biol. Chem. |
| TCEP | Sigma | Cat# C4706 |
| Dithiothreitol (DTT) | Roche | Cat# 10197777001 |
| Ni-NTA resin | Qiagen | Cat# 30210 |
| Casamino acids | Gibco | Cat# 223050 |
| cOmplete His-Tag purification beads | Roche | Cat# 5893682001 |
| EcoRI-HF Restriction Enzyme | New England Biolabs | Cat# R3101S |
| SYBR Safe DNA Gel Stain | Invitrogen | S33102 |
| TaqMan Fast Universal PCR Master Mix (2X) | Thermo Fisher | Cat# 4352042 |
| Apex General Purpose Agarose | Genesee Scientific | Cat# 20-102GP |
| Superscript III First-Strand Synthesis System | Thermo Fisher | Cat# 18080051 |
| 2X Universal SYBR Green Fast qPCR Mix | Abclonal | Cat# RK21203 |
| Oligonucleotides | | |
| MSSS-FWD (early): AACCCACTGCTTAAGCCTCA | Christensen et al. | N/A |
| MSSS-REV (early): ACCAGAGTCACACAACAGACG | Christensen et al. | N/A |
| FST-FWD (intermediate): AGCCGCCTAGCATTTCATCA | Christensen et al. | N/A |
| FST-REV (intermediate): CCAGCGGAAAGTCCCTTGTA | Christensen et al. | N/A |
| Late RT-FWD: TGTGTGCCCGTCTGTTGTGT | Christensen et al. | N/A |
| Late RT-REV: CTTCAGCAAGCCGAGTCCTG | Christensen et al. | N/A |
| cGAS gRNA target: GAGGCCGCCCTGCCTAAGGC | Johnson et al. | N/A |
| STING gRNA target: CCCCGTGACCCCTGGGACAC | Johnson et al. | N/A |
| IRF3 gRNA target: CGTGCGGCTCTTGTTCACCC | Johnson et al. | N/A |
| PQBP1 KO1 gRNA target: GTCTGCAGCGCAACGGGCAG | This study | N/A |
| PQBP1 KO2 gRNA target: GGATGCCTCTCTTGGCCAAG | This study | N/A |
| FST-FWD (HIV-1-GFP, intermediate): AGCCTCCTAGCATTTCGTCAC | This study | N/A |
| FST-REV (HIV-1-GFP, intermediate): CCAGCGGAAAGTCCCTTGTA | This study | N/A |
| TaqMan eGFP assay | Thermo Fisher | Cat# Mr04329676_mr |
| TaqMan IFITM1 assay | Thermo Fisher | Cat# Hs00705137_s1 |
| TaqMan IFITM3 assay | Thermo Fisher | Cat# Hs03057129_s1 |
| TaqMan MX1 assay | Thermo Fisher | Cat# Hs00895608_m1 |
| TaqMan ISG15 assay | Thermo Fisher | Cat# Hs01921425_s1 |
| TaqMan IFI27 assay | Thermo Fisher | Cat# Hs01086373_g1 |
| TaqMan GAPDH assay | Thermo Fisher | Cat# Hs02786624_g1 |
| Recombinant DNA | | |

| | | |
|---|---|---|
| psPAX2 | Gift from Didier Trono | Addgene plasmid #12260 |
| pCMV-VSV-G | Gift from Bob Weinberg | Addgene plasmid #8454 |
| pCA528 | Gift from Wes Sundquist | DNASU plasmid pCA528 |
| pCA528-huPQBP1 | This study | Addgene plasmid #217449 |
| lentiCRISPR v2 | Gift from Feng Zhang | Addgene plasmid #52961 |
| lentiCRISPR control (LCV2) empty vector minus stuffer | Johnson et al., 2018 | Addgene plasmid #217443 |
| lentiCRISPR cGAS | Johnson et al., 2018 | Addgene plasmid #217444 |
| lentiCRISPR STING | Johnson et al., 2018 | Addgene plasmid #217445 |
| lentiCRISPR IRF3 | Johnson et al., 2018 | Addgene plasmid #217446 |
| lentiCRISPR PQBP1 KO1 | This study | Addgene plasmid #217447 |
| lentiCRISPR PQBP1 KO2 | This study | Addgene plasmid #217448 |
| Software | | |
| GraphPad Prism 9.0 | GraphPad | https://www.graphpad.com/scientific-software/prism/ |
| FlowJo 10.0 | FlowJo LLC | https://www.flowjo.com/ |
| Image Studio Lite 5.5 | LI-COR | https://www.licor.com/bio/image-studio-lite/ |
| Morpheus | Broad Institute | https://software.broadinstitute.org/morpheus/ |
| Illustrator | Adobe | https://www.adobe.com/products/illustrator |
| BioRender | BioRender | https://www.biorender.com/ |
| Other Reagents | | |
| LS Columns | Miltenyi Biotec, Inc | 130-042-401 |
| CD14 MicroBeads, human | Miltenyi Biotec, Inc | 130-050-201 |
| QuadroMACS Separator | Miltenyi Biotec, Inc | 130-090-976 |
| ProFlex PCR System | Thermo Fisher | Cat# A41182 |
| QuantStudio 3 Real-Time PCR System | Applied Biosystems | A28137 |
| Attune NxT Flow Cytometer | Thermo Fisher | A24858 |
| Attune NxT Autosampler | Thermo Fisher | 4473928 |
| Odyssey CLx Imager | LI-COR | CLX-2063 |
| SpectraMax ID.5 Plate Reader | Molecular Devices | Cat #76175-288 |
| BioTek Synergy Neo2 Plate Reader | Agilent | BTNEO2 |
| SW 32 Ti Rotor, Swinging Bucket | Beckman | Cat# 369694 |
| 0.45µm syringe filters | Corning | Cat# 28200-026 |
| Optima Ultracentrifuge | Beckman | Model: LE-80k |
| Bolt 4-12% Bis-Tris Plus Gels | Invitrogen | NW04125BOX |
| HiTrap Heparin HP 5 mL | Cytiva | Cat #17040701 |
| HiTrap Q HP 5 mL | Cytiva | Cat #17115401 |
| Superdex 75, 120 mL;16/600 | Cytiva | Cat# 28989333 |
| Thinwall Polyallomer, Konical Tubes; 25 x 89 mm | Beckman | Cat# 358126 |
| Open-Top Thinwall Ultra-Clear Tube, 25 x 89 mm | Beckman | Cat# 344058 |
| 2',3'-cGAMP ELISA Kit | Arbor Assays | Cat# K067-H5 |
| LIVE/DEAD® Fixable Violet Dead Cell Stain Kit | Molecular Probes | L34955 |
| BCA protein assay, reducing agent compatible | Thermo Fisher | Cat# PI23252 |
| p24 ELISA | Xpress Bio | XB-1000 |
| PureLink HiPure Plasmid Maxiprep Kit | Invitrogen | K210007 |
| QIAquick PCR Purification Kit | Qiagen | Cat# 28106 |
| Cytofix/Cytoperm Fixation/Permeabilization Kit | BD Biosciences | 554714 |
| Human leukocytes from normal donors (deidentified) | ARUP Components | N/A |

| | | |
|---|---|---|
| DNeasy Blood & Tissue Kit | Qiagen | Cat# 69504 |
| QIAquick Gel Extraction Kit | Qiagen | Cat# 28706 |
| RNeasy 96 Kit | Qiagen | Cat# 74181 |
| Owl EasyCast B1 Mini Gel Electrophoresis System | Thermo Fisher | Cat# B1-BP |
| PowerEase Touch 250W Power Supply | Thermo Fisher | Cat# PSC350M |
| iBright FL1500 Imaging System | Thermo Fisher | Cat# A44115 |

**S1 Fig.**

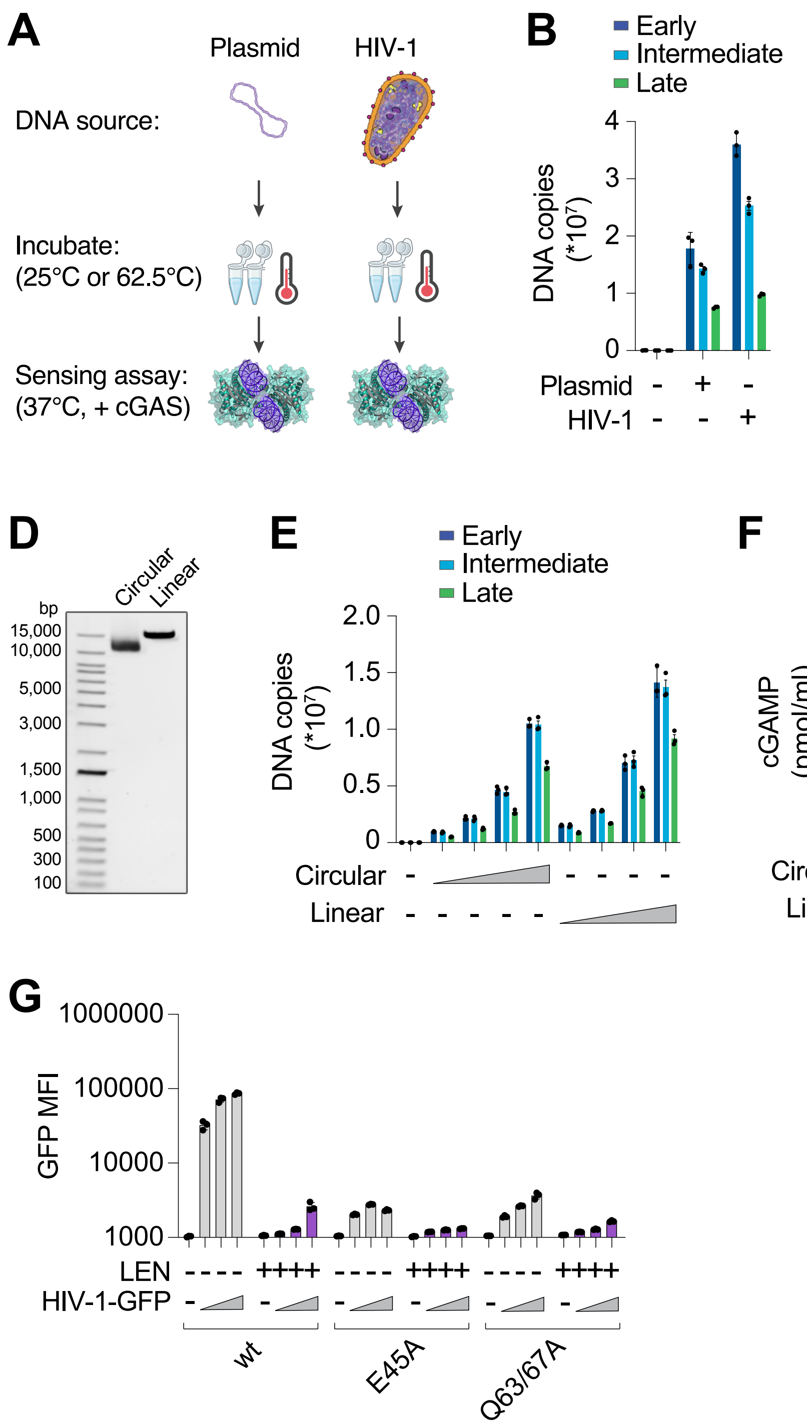
A
Plasmid
HIV-1
DNA source:
Incubate:
(25°C or 62.5°C)
Sensing assay:
(37°C, + cGAS)
B
Early
Intermediate
Late
DNA copies (*10^7)
Plasmid - + -
HIV-1 - - +
C
cGAMP (pmol/ml)
*
****
Plasmid - + + - -
HIV-1 - - - + +
Heat - - + - +
D
Circular
Linear
bp
15,000
10,000
5,000
3,000
1,500
1,000
500
300
100
E
Early
Intermediate
Late
DNA copies (*10^7)
Circular
Linear
F
cGAMP (pmol/ml)
Circular
Linear
G
GFP MFI
LEN
HIV-1-GFP
wt
E45A
Q63/67A

**S1 Fig. Capsid stability can be affected by heat or CA mutations.**

**(A)** Schematic depicting heat treatment of plasmid or ERT samples prior to cell-free cGAS sensing assays.

**(B)** Copy number of plasmid DNA or viral DNA from ERT reactions that were incubated under identical "standard" conditions (16 h at 37°C) to be taken through limited heat treatment.

**(C)** Cell-free sensing assay (related to panels A&B) comparing the ability of plasmid DNA and HIV-1 ERT samples to activate recombinant cGAS. After ERT, samples were incubated briefly (20 s) at 62.5ºC before assaying with cGAS under standard conditions (8 h for 37°C). Heat treatment slightly decreased the ability of plasmid DNA to activate cGAS (possibly through partial denaturation and/or aggregation of the DNA into a less accessible structure). In contrast, heat treatment of ERT products robustly increased cGAS activity.

**(D)** Gel electrophoresis image depicting a molecular weight ladder (lane 1), circular control DNA (pSG3Δenv plasmid, lane 2), and linear control DNA (pSG3Δenv plasmid digested with EcoRI, lane 3).

**(E)** qPCR measurements of DNA copy numbers from circular and linear control DNA samples from (D) that were incubated under standard ERT conditions (16 h at 37°C) in a 2-fold dilution series.

**(F)** Cell-free sensing assay (related to panels D&E) of circular and linear control DNA. Samples were incubated with recombinant cGAS and cGAMP levels were measured by ELISA.

**(G)** GFP mean fluorescence intensity measured by flow cytometry in THP-1 monocytic cells that were infected for 48 h with HIV-1-GFP, the capsid stabilized CA mutant E45A, or the destabilized mutant Q63/67A at matched virus doses (12.5, 25, and 50 nM p24). Where indicated, LEN (10 nM) was added at the time of infection.

For (C), statistics were calculated using an unpaired t test to compare plasmid or HIV-1 samples: $p<0.05$: *, $p<0.0001$: ****. Graphs show mean ± SD from three samples from a representative experiment, selected from two independent experiments.

S2 Fig.

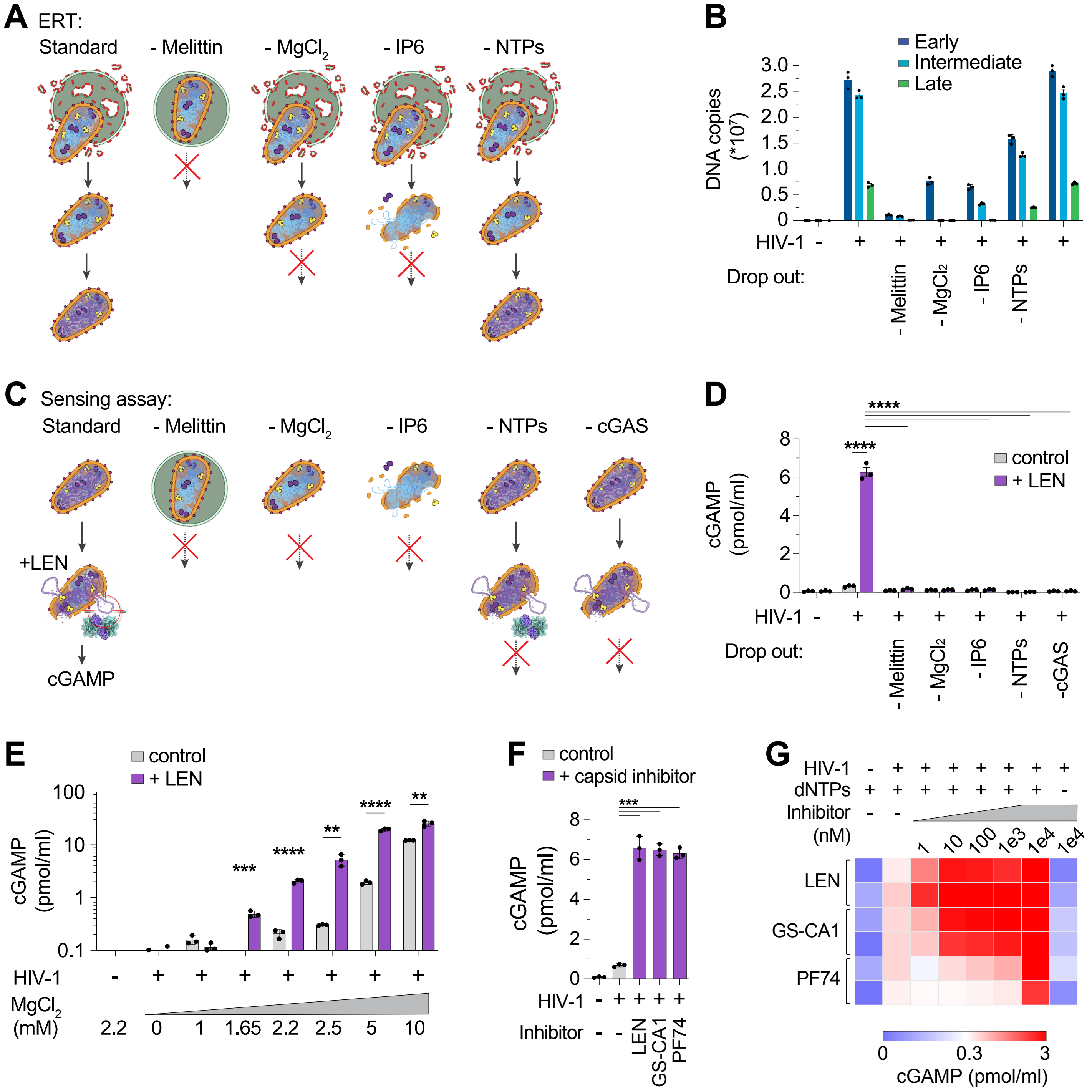
A
ERT:
Standard
- Melittin
- MgCl2
- IP6
- NTPs
B
Early
Intermediate
Late
DNA copies (*10^7)
HIV-1
Drop out:
- Melittin
- MgCl2
- IP6
- NTPs
C
Sensing assay:
Standard
- Melittin
- MgCl2
- IP6
- NTPs
- cGAS
+LEN
cGAMP
D
cGAMP (pmol/ml)
control
+ LEN
HIV-1
Drop out:
- Melittin
- MgCl2
- IP6
- NTPs
-cGAS
E
control
+ LEN
cGAMP (pmol/ml)
HIV-1
MgCl2 (mM)
2.2 0 1 1.65 2.2 2.5 5 10
F
control
+ capsid inhibitor
cGAMP (pmol/ml)
HIV-1
Inhibitor
LEN
GS-CA1
PF74
G
HIV-1
dNTPs
Inhibitor (nM)
1 10 100 1e3 1e4 1e4
LEN
GS-CA1
PF74
0 0.3 3
cGAMP (pmol/ml)

**S2 Fig. Cell-free sensing of HIV-1 requires virion permeabilization, magnesium, NTPs, and cGAS.**

**(A)** Schematic illustrating that under standard ERT reaction conditions, viral RNA is converted into DNA inside an intact or largely intact capsid. When key components are omitted from the reaction, the ERT reaction is inhibited at the indicated steps.

**(B)** qPCR measurements of DNA copy numbers from ERT reactions that were carried out for 16 h at 37°C under standard conditions (including melittin, $MgCl_2$, IP6, NTPs, dNTPs, and additional buffer components as stated in the Methods) compared to reactions performed in the absence of melittin, $MgCl_2$, IP6, or NTPs.

**(C)** Models for cell-free sensing reactions corresponding to ERT conditions depicted in (A). Core disruption with LEN permits cGAS sensing of RT products under standard reactions conditions. Sensing is blocked at the indicated steps when specific reaction components are removed.

**(D)** Cell-free cGAS sensing assay performed with ERT samples shown in (B). After ERT, samples were incubated with (or without) recombinant cGAS under the conditions shown, with or without LEN (100 nM). cGAMP levels were determined by ELISA.

**(E)** Cell-free sensing assay showing sensitivity to $MgCl_2$ concentration. Standard HIV-1 ERT samples were aliquoted into cGAS reactions with the indicated amounts of $MgCl_2$, and with or without LEN (100 nM).

**(F)** Cell-free sensing assay depicting core disruption by other capsid inhibitors. ERT samples that were prepared under standard conditions (16 h at 37°C) were aliquoted into cGAS reactions and incubated in the presence or absence of LEN (100 nM), GS-CA1 (100 nM), or PF74 (10 μM).

**(G)** Heat map showing dose-dependent increase in cGAS activity for samples incubated with LEN, GS-CA1, or PF74. Samples were prepared for cGAS assays as in (F) with increasing amounts of the indicated capsid inhibitors. Two replicates shown from one of three independent experiments. Statistics were calculated using a one-way ANOVA with Tukey's multiple comparisons test in (D,F) or an unpaired t test for each concentration of $MgCl_2$ in (E): $p<0.01$: **, $p<0.001$: ***, $p<0.0001$: ****. Graphs depict mean ± SD from three samples from a representative experiment, selected from three independent experiments.

**S3 Fig.**

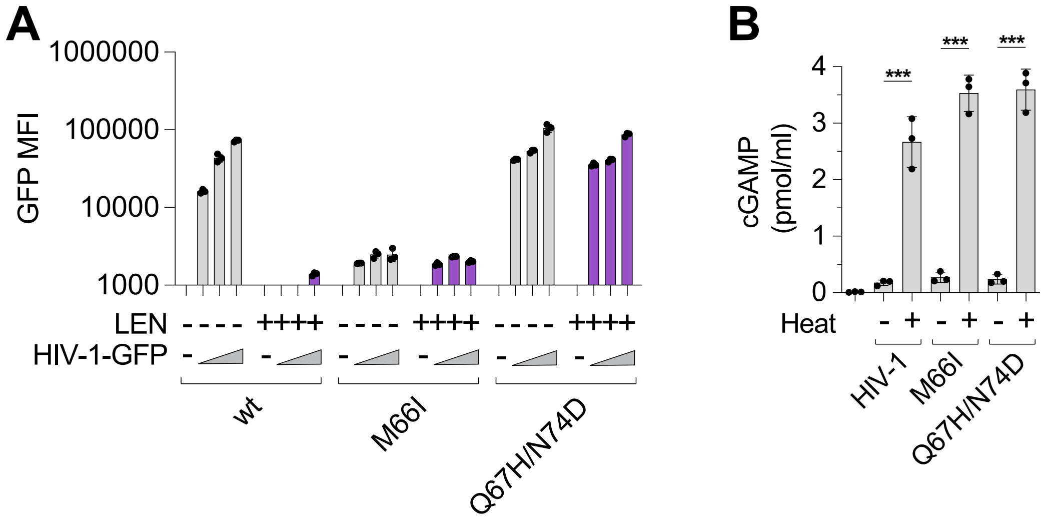

A
1000000
100000
10000
1000
GFP MFI
LEN
HIV-1-GFP
wt
M66I
Q67H/N74D
B
cGAMP (pmol/ml)
4
3
2
1
0
***
Heat
HIV-1
M66I
Q67H/N74D

**S3 Fig. Specific CA mutations confer resistance to LEN.**

**(A)** Mean fluorescence intensity of GFP measured by flow cytometry in THP-1 monocytic cells infected for 48 h with HIV-1-GFP compared to LEN-resistant CA mutants M66I and Q67H/N74D at matched virus doses (12.5, 25, and 50 nM p24). Where indicated, LEN (10 nM) was added at the time of infection.

**(B)** Cell-free sensing assay of wt, M66I, Q67H/N74D virions. Samples from ERT reactions shown in Fig. 3A that were incubated under standard conditions (16 h of ERT at 37°C) were then briefly heat treated (20 s at 62.5°C) before incubating with recombinant cGAS for 8 h at 37°C. cGAMP levels were measured by ELISA. Statistics were calculated using a one-way ANOVA with Tukey's multiple comparisons test: $p<0.001$: ***. Graphs depict mean ± SD from three samples from a representative experiment, selected from 2 (A) or 3 (B) independent experiments.

**S4 Fig.**

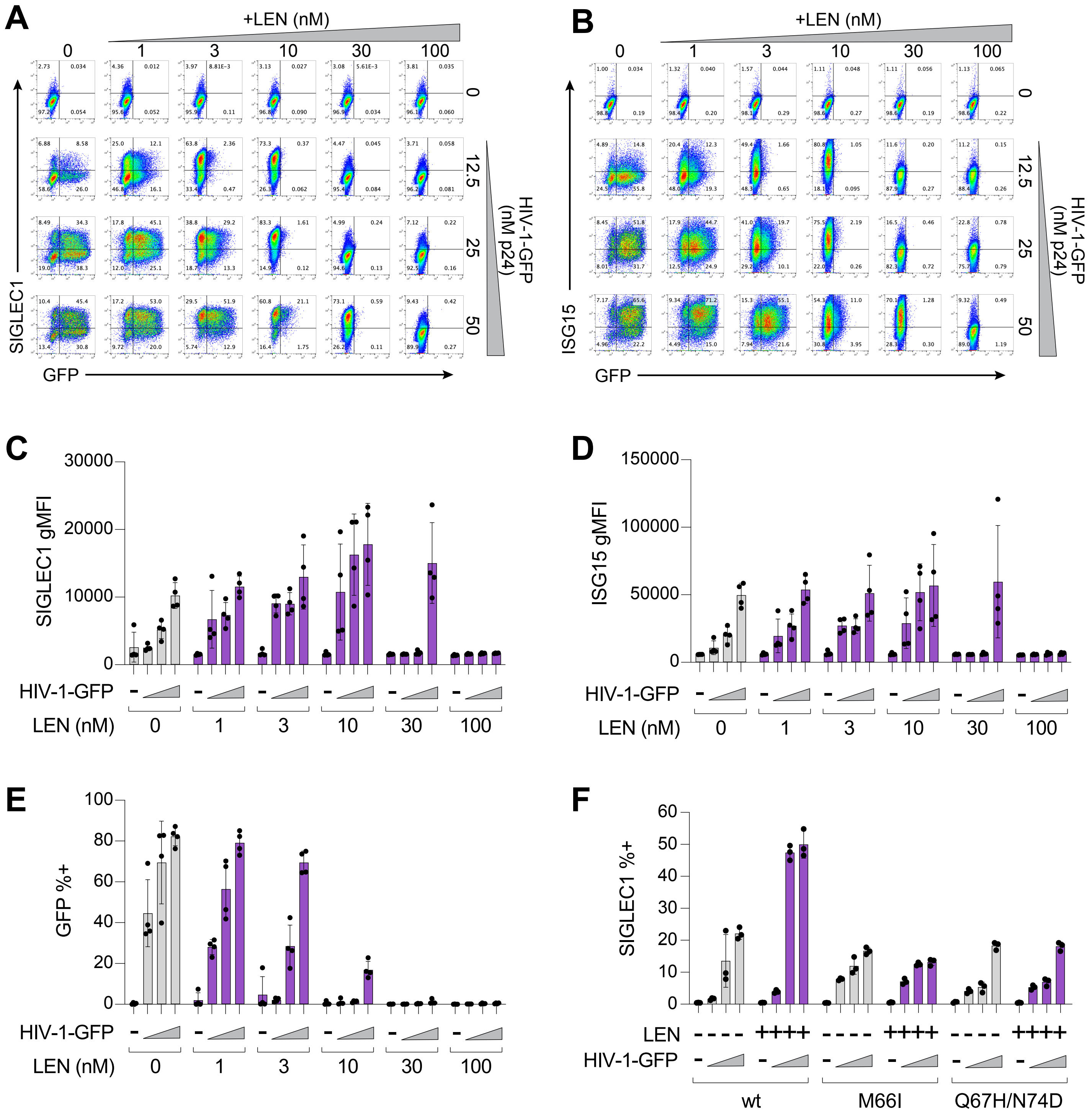
A
+LEN (nM)
0
1
3
10
30
100
12.5
25
50
HIV-1-GFP (nM p24)
SIGLEC1
GFP
B
+LEN (nM)
ISG15
GFP
HIV-1-GFP (nM p24)
C
SIGLEC1 gMFI
30000
20000
10000
0
HIV-1-GFP
LEN (nM)
D
ISG15 gMFI
150000
100000
50000
0
HIV-1-GFP
LEN (nM)
E
GFP %+
100
80
60
40
20
0
HIV-1-GFP
LEN (nM)
F
SIGLEC1 %+
60
50
40
30
20
10
0
LEN
HIV-1-GFP
wt
M66I
Q67H/N74D

**S4 Fig. LEN augments innate immune responses across a range of virus doses.**

**(A-B)** Flow cytometry of THP-1 monocytic cells infected for 48 h with a range of HIV-1-GFP doses (12.5, 25, and 50 nM p24) and increasing amounts of LEN. GFP is displayed on the x-axis and either SIGLEC1 (A) or ISG15 (B) are shown on the y-axis. Plots represent one of four infection replicates, which are averaged in the heat maps shown in Fig. 4C.

**(C-E)** Flow cytometry data from THP-1 cells infected with a range of HIV-1-GFP doses (12.5, 25, and 50 nM p24) for 48 h and increasing amounts of LEN (0, 1, 3, 10, 30, and 100 nM), showing SIGLEC1 gMFI (C), ISG15 gMFI (D), or %GFP+ (E).

**(F)** Percent SIGLEC1-positive cells measured by flow cytometry. THP-1 monocytic cells were infected for 48 h with HIV-1-GFP compared to LEN-resistant CA mutants M66I and Q67H/N74D at matched doses (12.5, 25, and 50 nM p24). LEN (10 nM) was added at the time of infection. Graphs show mean ± SD from three-four samples from a representative experiment, selected from three (C-E) or two (F) independent experiments.

**S5 Fig.**

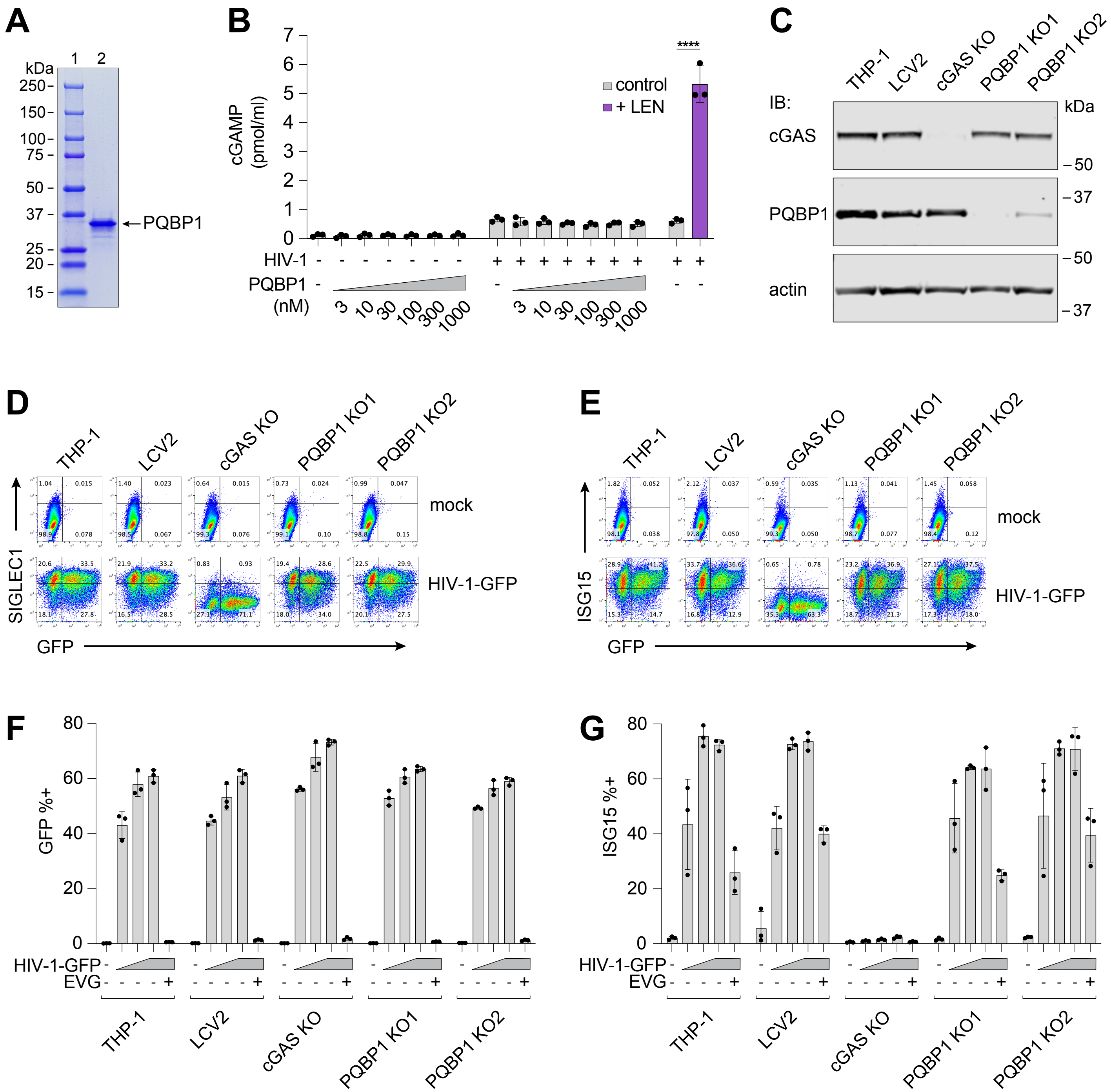

A
kDa
1
2
250
150
100
75
50
37
25
20
15
PQBP1
B
cGAMP (pmol/ml)
control
+ LEN
****
HIV-1
PQBP1 (nM)
3
10
30
100
300
1000
C
THP-1
LCV2
cGAS KO
PQBP1 KO1
PQBP1 KO2
IB:
cGAS
PQBP1
actin
kDa
50
37
D
THP-1
LCV2
cGAS KO
PQBP1 KO1
PQBP1 KO2
mock
HIV-1-GFP
SIGLEC1
GFP
E
THP-1
LCV2
cGAS KO
PQBP1 KO1
PQBP1 KO2
mock
HIV-1-GFP
ISG15
GFP
F
GFP %+
HIV-1-GFP
EVG
THP-1
LCV2
cGAS KO
PQBP1 KO1
PQBP1 KO2
G
ISG15 %+
HIV-1-GFP
EVG
THP-1
LCV2
cGAS KO
PQBP1 KO1
PQBP1 KO2

**S5 Fig. PQBP1 does not affect cGAS activity *in vitro* and is not essential in cells.**

**(A)** SDS-PAGE with Coomassie staining showing a molecular weight ladder (lane 1) and purified recombinant human PQBP1 protein (lane 2).

**(B)** Cell-free sensing assay to determine cGAS activity in the presence or absence of PQBP1. Purified PQBP1 was incubated at increasing concentrations with recombinant cGAS alone, or together with HIV-1 samples aliquoted from standard ERT reactions (16 h at 37°C). Cell-free sensing assay reactions were performed under standard conditions. LEN (100 nM) was included as a positive control for viral capsid disruption.

**(C)** Immunoblots of THP-1 lysates after lentiCRISPR editing, depicting KO of cGAS or PQBP1 (two independent gRNAs targeting PQBP1), relative to a non-targeting vector control (LCV2).

**(D)** Flow cytometry of lentiCRISPR-modified THP-1 cells infected with HIV-1-GFP for 48 h (50 nM p24), depicting GFP expression on the x-axis and SIGLEC1 on the y-axis.

**(E)** Flow cytometry of lentiCRISPR-modified THP-1 cells infected with HIV-1-GFP for 48 h (50 nM p24), depicting GFP expression on the x-axis and ISG15 on the y-axis.

**(F-G)** Flow cytometry data from THP-1 cells infected with a range of HIV-1-GFP doses (12.5, 25, and 50 nM p24) for 48 h, showing %GFP+ (F) or %ISG15+ (G). To evaluate whether cGAS or PQBP1 have roles in sensing early steps in the virus life cycle, we also tested HIV-1-GFP (50 nM p24) in the presence of the integration inhibitor elvitegravir (EVG, 1 μM). All graphs show mean ± SD from three samples from a representative experiment, selected from two independent experiments.
